# Delta variant with P681R critical mutation revealed by ultra-large atomic-scale *ab initio* simulation: Implications for the fundamentals of biomolecular interactions

**DOI:** 10.1101/2021.12.01.470802

**Authors:** Puja Adhikari, Bahaa Jawad, Praveen Rao, Rudolf Podgornik, Wai-Yim Ching

## Abstract

SARS-CoV-2 Delta variant is emerging as a globally dominant strain. Its rapid spread and high infection rate are attributed to a mutation in the spike protein of SARS-CoV-2 allowing the virus to invade human cells much faster and with increased efficiency. Particularly, an especially dangerous mutation P681R close to the furin cleavage site has been identified as responsible for increasing the infection rate. Together with the earlier reported mutation D614G in the same domain, it offers an excellent instance to investigate the nature of mutations and how they affect the interatomic interactions in the spike protein. Here, using ultra large-scale *ab initio* computational modeling, we study the P681R and D614G mutations in the SD2-FP domain including the effect of double mutation and compare the results with the wild type. We have recently developed a method of calculating the amino acid-amino acid bond pairs (AABP) to quantitatively characterize the details of the interatomic interactions, enabling us to explain the nature of mutation at the atomic resolution. Our most significant find is that the mutations reduce the AABP value, implying a reduced bonding cohesion between interacting residues and increasing the flexibility of these amino acids to cause the damage. The possibility of using this unique mutation quantifiers in a machine learning protocol could lead to the prediction of emerging mutations.

## 1. INTRODUCTION

COVID-19 pandemic started two years ago and continues with unabated intensity with no clear end in sight, despite many repeated attempts to contain it. The recent emergence of various variants of concern (VOC) in Severe Acute Respiratory Syndrome Coronavirus-2 (SARS-CoV-2) ^1^ like Alpha ^2^, Beta ^3^, Delta ^4^, and Gamma ^5^, and variants of interest (VOI) such as Eta ^6^, Iota ^7^, Kappa ^8^, Lambda ^9^, and Mu ^10^, instigates new anxieties. Among the new VOC, the Delta variant causes more severe infection and spreads faster than previous variants of the SARS-CoV-2 virus, emerging as the dominant strain in the world ^11^, causing worries among the general population and solidifying the belief that the battle against the pandemic will be a long one. In a broader context, this historical moment that faces us is a grand one from every conceivable direction. It also introduces a new chapter in the perception and significance of biomedical sciences. However, a successful response to this dire situation crucially implies and promotes not only pandemics related efforts in biomedical sciences but even more importantly deep level collaborations across all scientific fields guiding a concerted action grounded in different social instruments.

The evolution of viruses in recent decades has been well-documented, including the 1918 flu pandemic ^12^, the zoonotic HIV ^13^, the seasonal flu virus variations as well as several predecessors of SARS-CoV-2 such as SARS-CoV-1 in 2003 and MERS in 2012 ^14^. The emergence of the Delta variant together with other VOC are a *natural and unavoidable part of the virus evolution*, and can be traced to specific mutations of the amino acids in the protein sequence that can result in an enhanced infection rate or can quench the full action of the already developed vaccines and/or other therapeutic drugs ^15, 16^. Other mutational variants, in addition to the known Alpha, Beta, Gamma, Delta etc., will continue to emerge as the epidemic rages on, making it imperative to strive for a fundamental understanding of the role of mutations at a deeper molecular and atomic level. This fundamental understanding can enable the design of new strategies and methods to combat the current and the emerging variants such as the AY.4.2 “Delta plus” variant, that seems to be more transmissible than the original Delta variant in the United Kingdom ^17^.

The spike (S) protein of SARS-CoV-2 has two subunits, S1 and S2, responsible for ACE2 receptor docking and membrane fusion, respectively ^18^. In fact, SARS-CoV-2 enters the host cells through its S-protein, which is synthesized as an inactive precursor that must be cleaved to successfully mediate membrane fusion ^19^. The cleavage activation mechanism occurs at S1/S2 and S2' cleavage sites ^20^, the former being located at the boundary between the S1 and S2 subunits having a unique polybasic insertion furin recognition site _681_PRRAR|S_686_ (| denotes proteolytic cleavage site) ^19^. The S-protein is thus first cleaved at the S1/S2 site, which does not actually result in the complete separation of the S1 and S2 subunits but allows them to remain non-covalently bound ^20^. Upon the S1/S2 cleavage and binding of S-protein to ACE2, a second cleavage site, S2', becomes more exposed to be completely cleaved by host proteases for activating virus-cell membrane fusion ^19-25^. Hence, the unique S1/S2 site has been identified as mainly responsible for its high infectivity and transmissibility ^22^. Interestingly, the P681R mutation right at the furin cleavage site of the Delta variant plays an important role in enhancing the S-protein cleavage ^26-28^ and is hypothesized as the main culprit for the Delta variant infectivity ^26-28^. In addition to P681R mutation, the Delta variant contains also D614G mutation that promotes the RBD of S-protein in an “open” conformation, making its binding with the ACE2 receptor easier ^29^, as well as enhancing the protease cleavage at the S1/S2 cleavage site ^30^. In view of this overarching importance of the P681R and D614G mutations, it is therefore crucial to understand the role that these mutations play in the phenomenology of the Delta variant at the *atomic scale*, which can only be accomplished by unleashing the best that the *ab initio* quantum chemical methodology has to offer.

The specific aim of this study is to investigate the nature of these two important mutations, P681R and D614G in the Delta variant using ultra large-scale *ab initio* quantum chemical modeling, combined with advanced analysis that allows for a quantitative assessment of the impact of mutations on the atomic resolution scale. Specifically, we study atomically resolved structure and quantify the interatomic impact of P681R and D614G in the SD2 to FP (SD2-FP) domains of the S-protein together with the effect of the double mutation and compare the results with the unmutated case or the wild type (WT). We use the recently developed method of calculating the amino acid-amino acid bond pairs (AABP) to characterize the quantitative details of the interatomic interactions ^31^. Such unprecedented and detailed analysis of the origin and impact of atomistically resolved mutations provides many fundamental insights that could lead to a new level of understanding in the development of therapeutic drug design against the SARS-CoV-2 virus and its variants.

## 2. MODEL DESIGN AND CONSTRUCTION

Large biomolecular systems such as proteins have complex structures and contain many amino acids linked together in a specific order. Currently, we are capable to conduct *ab initio* simulations with up to 5000 atoms for a single calculation by adopting a *divide and conquer* strategy to investigate the S-protein. We briefly describe the model used in this study.

The SD2 to FP region marked as region 3 in **Figure S1** is our SD2-FP model which is used in the actual atomic-scale calculation in the present work. The other broader structural regions consist of different structural domains in the S-protein of the SARS-CoV-2 which shows the specific mutation sites in the Delta variants. This is fully illustrated in **Figure S1** in the Supplementary Information (**SI**). The SD2-FP models involved in the present work are: (a) the wild type (WT), (b) the mutated model P681R, (c) the mutated model D614G and (d) the double mutation with both D614G and P681R.

The initial structure for the regions shown in **Figure S1** was obtained from Woo *et al* from [PDB ID 6VSB] ^32^ which originated from Wrapp et al. study ^18^. Here it should be mentioned that the 6VSB Cryo-EM structure from Wrapp et al. has many missing amino acids (AAs) due to the limitation in their technique. Different prediction methods to model the missing AAs of 6VSB were used and the details of obtaining the full-length SARS-CoV-2 spike (S) protein structure can be found in Woo *et al* ^32^. We chose Chain A in its up conformation. Sequence numbers 592-834 in S-protein were used for SD2-FP model (6VSB_1_2_1) ^33^. We used our procedure to construct a manageable size model and summarize it as follows. First, we selected all residues of the SD2 and FP region to create the SD2-FP model (residue 592 to 834). Next, we removed the glycans and the associated hydrogen (H) atoms from the SD2-FP model and later added the H atoms using the Leap module with ff14SB force field in the AMBER package ^34-36^, which then yields the WT model used as a template to generate the mutated models. To prepare the mutated models with a single mutation (P681R or D614G) or a double mutation (P681R and D614G), we used Dunbrack backbone-dependent rotamer library ^37^, implemented by the UCSF Chimera package ^38^. In total, we have created four SD2-FP models, including the WT model, mutated with P681R and D614G mutations, and double mutation consisting of both D614G and P681R. These models were minimized with 100 steepest descent steps and 10 conjugate gradient steps using UCSF Chimera to avoid bad clashes. These models (see **Figure 1** and **S1**) were then further optimized using Vienna *ab initio* simulation package (VASP) ^39^ as described in **subsection S1.1** in **SI**.

**Figure 1.**
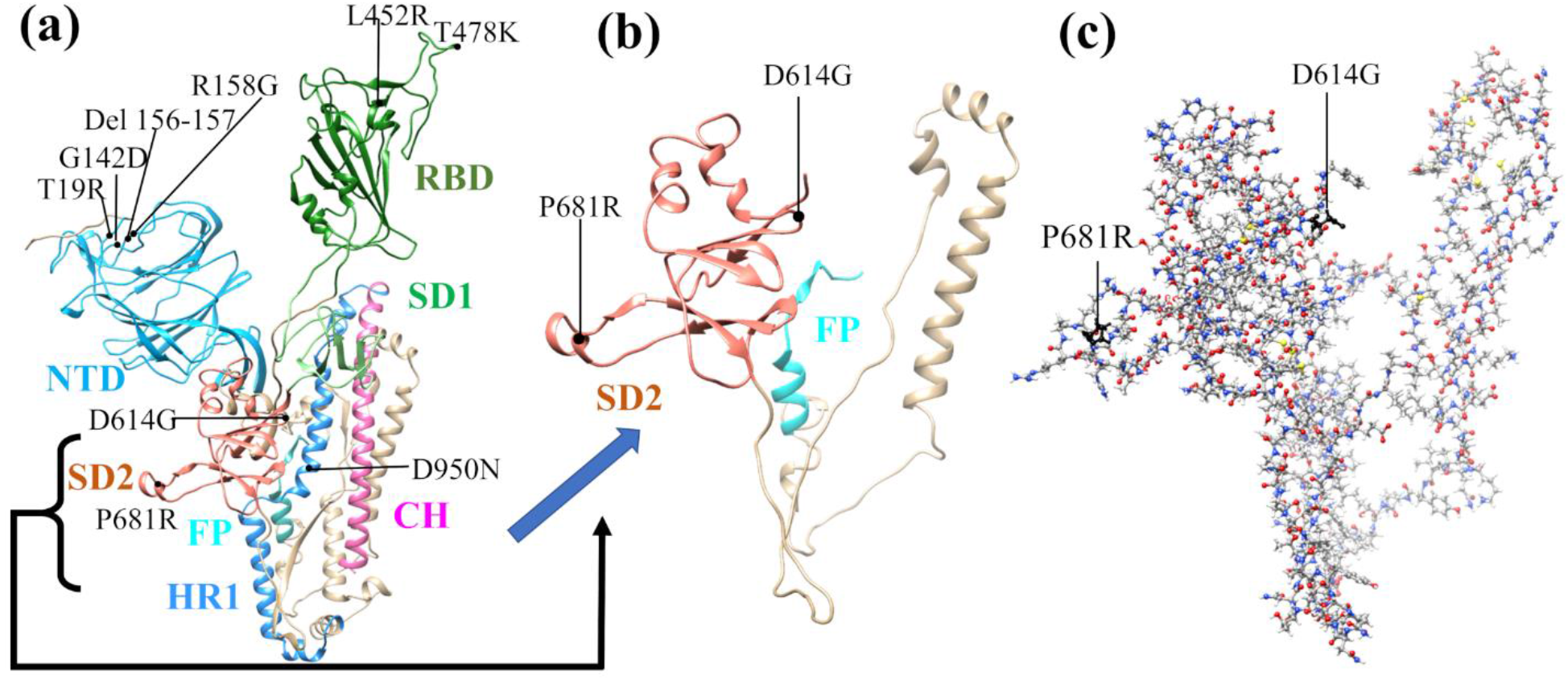
The illustration of SD2-FP model construction. **(a)** Ribbon of a single protomer in up conformation (chain A) of spike protein SARS-CoV-2 from signal peptide (SP) to central helix (CH) and the associated mutation for Delta variant in different domains, **(b)** the ribbon structure of SD1-FP model with two marked mutations that is selected for constructing our models, **(c)** ball and stick of the SD1-FP in (b) and their respective mutations as marked.

## 3. AMINO ACID – AMINO ACID BOND PAIR (AABP)

The VASP optimized structures were used as input in *orthogonalized linear combination of atomic orbitals* (OLCAO) method ^40^ to calculate the electronic structure and interatomic interactions in biomolecules. The details of OLCAO method are described in **section S1.2** of **SI**. Using OLCAO we calculated the *bond order* (BO), which quantifies strength of the bond between two atoms and usually scales with *bond length* (BL). The sum of all BO values within a single structure unit gives the *total bond order* (TBO). The relatively new concept of BO and TBO in biomolecules quantifies the cohesion of the system. Before we extend our formulation and analysis of BO and TBO to the *amino acid - amino acid bond pair* (AABP), we will first discuss briefly the 20 canonical amino acids with distinct residues listed in **Table S1** and illustrated through their functional groups in **Figure S2**.

Amino acids are the basic structural units of proteins, sharing three common structural elements: an amine group, a carboxyl group, and a side chain residue. Different functional groups comprising the side chain consign to each of the 20 canonical amino acids distinct physical properties that influence protein structure and function. The peptide bond links two adjacent AAs with a covalent bond between C1 (carbon number one) of one AA and N2 (nitrogen number two) of another, creating a linear chain connecting AAs with chemically distinct side chain residues into different linear sequences that can form long polypeptide chains, able to fold upon themselves and thereby giving rise to diverse, functionally distinct proteins. Any discussion of the structure, properties, and functionalities of proteins must therefore originate from the unique structures and properties of the 20 canonical AAs ^41, 42^.

Various physical properties characterize and differentiate the canonical AAs in bathing aqueous solutions, such as the number of atoms, size, molecular weight, hydropathy, polarity, charge, protonation/deprotonation dissociation constants, etc. In view of our aspiration to embark on a detailed *ab initio* investigation, unprecedented in size and scope, of the nature and effects of point mutations on interatomic interactions in the spike protein, most of the listed quantifiers do not seem to be appropriate and some appear to be distinctly missing. Based upon what the ultra-large scale *ab initio* methodology can provide, we focus on two structural quantifiers: the well-known *partial charge* (PC) parameter and a modification of the BO parameter referred to as the *amino acid - amino acid bond pair* (AABP). The first one addresses the polarity and charge structure of the molecule and certainly responds to all the alterations wrought by the mutation in the protein sequence. The second one is a generalization of the BO and TBO and is specifically tailored to proteins by defining the *amino acid - amino acid bond pair* (AABP) as

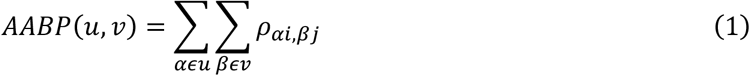

where the summations are over all atoms *α*in *AA u* and all atoms *β*in *AA v*. AABP embodies all possible bonding between two amino acids including both covalent and hydrogen bonding (HB). AABP is a single parameter that quantifies the amino acid-amino acid interaction, so that the stronger the interaction, the larger will be the AABP and vice versa.

We stress that the use of this novel AABP concept is not the same as using the conventional description of *amino acid-amino acid interaction* in biomolecules, since the distance of separation and the atomic interactions between two AAs are difficult to quantify accurately. The AABP values are calculated from quantum mechanical wave functions to study different types of interaction in biomolecules such as *nearest neighbor* (NN) and *non-NN*, also designated as *off-diagonal* or *non-local* (NL) interactions between AAs. Non-NN AAs are not vicinal in the 1D primary sequence space but are vicinal in the 3D embedding folding space. The AABP concept can help to foment a better understanding of the overall 3D interactions not only of proteins but of complex biomolecular systems in general ^31^.

The AABP defined above is a unique feature in the present study. Simulation methodologies that have been routinely and extensively used by the biomolecular research community ^43, 44^, such as classical molecular dynamics (MD) with its onerous energy or enthalpy calculations, are based on different types of presumably transferable *a-priori* force field specifications. As such they cannot reveal the atomic details of the real interatomic interactions and mostly rely on the atomic potential parametrizations and assumed geometric structures, which are both inherently limited. On the other hand, *ab initio molecular dynamics* methodology ^45^, intended for more realistic simulation of complex biomolecular systems and processes from first principles, is at present hampered by the excessive computational times and resources, making it inapplicable to the analysis of even modest size proteins. The use of the concept of bond order, as implemented in this work, can quantitatively characterize the AA-AA interaction in 3D folding space and can be applied also to larger scale protein-protein interaction, or the interaction of different segments of the same protein, thus providing a *promising and valuable alternative* (see section 4.2 for details).

Our approach here will be based on the characterization of the wild type and mutated protein by the PC and AA-AA bond pair parameters. By judiciously labeling each mutation as a data point, with specific details for the different components of the partial charge and AA-AA bond pair parameters, will furthermore facilitate its application in a machine learning (ML) protocol when many mutation data become available.

## 4. RESULTS

This section is conveniently divided into four subsections, but the key subsection is **4.3** AABP data for mutations in the structural domain of SD2-FP containing the furin cleavage site (**Figure S1**). The AABP data are based on the results of the model structures using VASP and the OLCAO for the electronic interactions. **Table S2** lists the structure information from VASP optimization for the four SD2-FP models in the Delta variant. (a) wild type (WT), (b) mutated P681R (R681), (c) mutated D614G (G614) and (d) double mutation (DM) labeled as G614-R681. In addition, **Table S2** also lists two HR1-CH models in the Delta variant. (e) wild type (WT) D950 and (f) mutated D950N (N950). As can be seen in **Table S2**, the energy is sufficiently converged to the level of 0.03 to 0.04 eV, less than 10^−5^ eV per atom including H atoms, but the entailed computational resources consumed are humongous. The VASP optimized structure is used as the input for the DFT calculation using OLCAO.

The results are divided into the following four subsections. Subsections **4.1** and **4.2** are standard electronic structures routinely present for the analysis in biomolecular systems ^31, 46-50^. **Subsection 4.2** describes the key data on interatomic interactions between all atoms whose results are used for the main **subsection 4.3** on the AABP data for the four SD2-FP models (a) to (d) listed in **Table S2. Section S2** in **SI** provides the additional results from the two HR1-CH models (e) and (f) listed in **Table S2** that support the observation in **subsection 4.3**.

### 4.1 Electronic structure

In *ab initio* calculation of any materials, the focus is on the density of states (DOS) or its components, the partial DOS (PDOS). In small molecules, researchers tend to use the list of energy levels separated by HOMO-LUMO gaps. **Figure S3** shows the calculated total DOS (TDOS) of the current supercell WT SD2-FP containing P681 & D614 with 3654 atoms. The 0.0 eV energy stands for the HOMO state or the top of the occupied valence band. The LUMO is located at about 2.5 eV. There exist some gap states within the HOMO-LUMO gap as expected due to some interacting states within this complex biomolecule. There is virtually no difference in the TDOS between the WT model and those that contain the mutated AA. The only difference is a very minute structure in some peaks, where presumably the mutated AA has a slightly different energy level (not shown). In principle, the PDOS can be resolved into individual AA or groups of AAs which will be very useful if a more detailed analysis is necessary such as making the distinctions between mutated and unmutated AAs. Atomic scale interaction must be revealed by detailed analysis of the calculated electronic structure on relevant AAs that will be fully revealed in this Section later.

**Figure S4** displays the PC on each of the 243 AAs from F592 to I834 in the WT SD2-FP model. The PC values of the mutated AAs G614, R681 and the double mutation G614-R681 are also highlighted and marked. The distribution of PC can be divided into three groups: (1) 20 AAs that are largely positively charged with PC values above 0.2 e^-^; (2) 21AAs largely negatively charged with PC values lower than -0.2 e^-^; and (3) a large group of 202 AAs with small PC between 0.2 e^-^ and -0.2 e^-^. The most positively charged AAs are W633 (2.01 e^-^) and Y764 (1.90 e^-^) and the majority of the highly negatively charged 13 AAs have nearly equal PC around -0.80 e^-^ to -0.99 e^-^. The data in **Figure S4** clearly show that the PC of D614 and P681 have dramatic changes in PC values under mutation and double mutation. D614 changes from the - 0.96 e^-^ in WT to -0.02 e^-^ and -0.01 e^-^ for mutated G614 and double mutation G614-R681, respectively. Similarly, P681 changes from 0.15 e^-^ for WT to 0.93 e^-^ and 1.03 e^-^ for mutated R681 and double mutation G614-R681. Besides the famous 614^th^ and 681^st^ residues there exists another site which shows high variation in PC: Y756. Y756 changes from a highly positive charge of 1.90 e^-^ in WT to -0.10 e^-^ in all mutated models.

**Figure S5** displays the PC of AAs on the solvent excluded surface of the SD1-FP model (**Fig.1 (c)**) with the location of key AAs that undergo mutation marked D614 and P681. Interestingly, D614 is highly negatively charged and P681 is near neutral (0.15 e^-^). Other relatively positively charged AAs (colored blue) shown in the **Figure S5** are F592, W633, R634, R646, R682, R683, R685, Y756, R765, K776, K786,

K790, and K814, whereas the negatively charged AAs (colored red) consist of E619, D627, E654, D663, E702, E725, D745, E748, E773, E780, D830, and I834. These results indicate that the atomic-scale calculations can provide charge distribution of protein subunits impacting the long-range electrostatic interaction between different structural units of the protein. The actual PC of selected AAs are listed in **Table S3**. PC distribution of HR1-CH WT model is displayed in **Figure S6**. The WT D950 has largely negative PC which changes to positive PC when mutates to N950. PC for each AAs for HR1-CH model is listed in **Table S4**.

### 4.2 Interatomic bonding

In contrast to the TDOS discussed above, the actual interatomic interaction in the form of bond order (BO) values (see **Computational Methods section S1** in **SI**) are fully available for all the atoms in the supercell used in the DFT calculation with the OLCAO method. These BO values are the basic ingredient of calculating the AABP values, central to this paper.

Figure 2. shows the BO vs BL distribution for every pair in the WT model for BL less than 3.5 Åincluding all covalently bonded pairs as well as the HB. The inset shows the distribution for BL from 2.0 Åto 4.5 Å. This is a very busy figure containing many interesting facts. We succinctly summarize them below:

1. The first group of covalent bonds are O-H, N-H and C-H with BL ranging from less than 1 Å to less than 1.2 Å with BO ranging from 0.15 e^-^ to 0.48 e^-^ depending on the actual structure of the AAs listed in **Figure. S2** and may even between certain AAs. There are two O-H bonds at 1.33 Å and 1.44 Å. The former bond occurs between two AAs (D808 and K811) and the latter one is from the same AA F592.
2. The second group of covalent bonds is the usual covalent bond between C, O and N. Their BO values can be very large ranging from 0.16 e^-^ for N-C up to 0.62 e^-^ for C=C which are strong double bonds. The relatively weaker C-C bonds have slightly larger BL. Similar observation can be seen for bonds between (C, O) and (N, C) pairs.
3. The next important bond is the HBs, mostly O⋯H and a few N⋯H. HBs are much weaker than covalent bonds but ubiquitous ranging all the way to ‘BL’ of close to 4 Å (see inset). According to a detailed analysis by Lei et al on a super-cold network of water ^51^, the maximum BO for O⋯H is around 0.1 e^-^.
4. The next interesting bonds are the covalent H-S and C-S bonds from the only S-containing AAs Cys and Met. Some H-S bonds have short BL (1.4 Å) with strong BO (0.3 e^-^) while the others have large BL with weak BO (more than 2.5 A and less than 0.05 e^-^). The C-S bonds are located at BL around 1.8 Å and with a relatively strong BO value between 0.1 e^-^ and 0.2 e^-^ at this longer BL. Our results show there are no disulfide bonds in the SD2-FP model.
5. We now focus our observations on the inset of **Figure 2** for BL ranging from 2.0 Åto 4.5 Å. It reveals many weaker HBs with BO less than 0.03 e^-^. Even more surprising is the presence of many atomic pairs (H-H, C-H, C-C, N⋯H) that contribute to BO values with weak but nontrivial BO values of less than 0.02 e^-^. These bonds are obviously formed between the non-local AAs which play a critical role in the total AABP values to be discussed in the next section.
6. One point we must emphasize is that the use of BO is a relatively new concept advocated by us. The BLs must be interpreted as the distance of separations between a pair of atoms, with the proviso that their interatomic interaction can go beyond the actual atomic pairs labeled as ‘BL’ due to the quantum effects arising from overlapping orbitals of their nearby atoms. Such subtle issues are usually ignored in biomolecular systems since they are seldom discussed in the context of quantum mechanical wave functions but rely on the distances between 2 atoms quantified by ‘BL’. Similar issues have been raised recently in the literature regarding the nature of C-H and C-C bonds ^52^.

**Figure 2.**
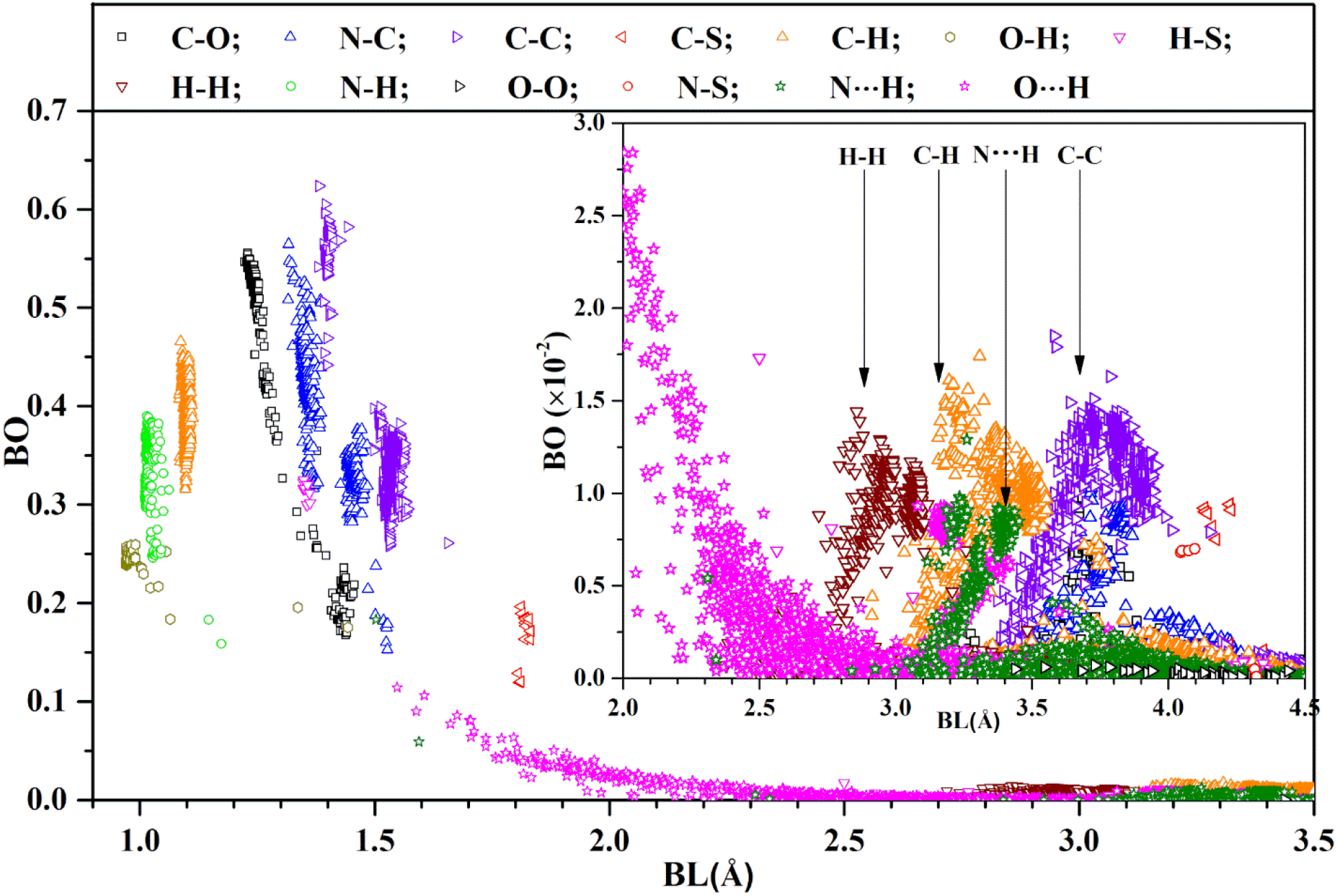
Distribution of BO vs BL for the WT SD2-FP model (1 - 3.5 Å). Inset: detailed distribution in range (2.0 Å-4.5 Å) with reduced BO scale. Most of them are in 4 types of atomic pairs marked with vertical arrows.

### 4.3 AABP data for mutations in Delta variant

The mutations on the Delta variant have been a hot topic that has attracted a lot of attention ^26, 28, 53-57^. Most of these studies focus on the clinical or experimental observations to demonstrate the danger of mutations especially the P681R near the furin cleavage site in the SD2-FP domain of the S-protein, but to the best of our knowledge, no theoretical explanation or computational studies have been reported so far. Based on the detailed atomic-scale electronic structure calculation described in the above two subsections, we extend the calculation of interactions between AAs involved in the mutations in the form of AABP described in the methods section. The calculated AABP values of mutations are summarized in **Table 1**. Each calculation is considered as a data point labeled by the specifically designed notation that will be instrumental in data mining and machine learning (ML) application (see **section 5.2**). The main observation of **Table 1** are as follows:

**Table 1.** AABP of four SD2-FP models and other information. DM denotes double mutation.

| Models | Total AABP | NN AABP | Non-local AABP | AABP from HB | No. of NL AAs | Data notation |
| --- | --- | --- | --- | --- | --- | --- |
| WT P681 | 1.117 | 1.064 | 0.054 | 0.064 | 5 | P681-1.117-1.064-0.054-0.064-0 |
| WT D614 | 0.917 | 0.912 | 0.005 | 0.040 | 5 | D614-0.917-0.912-0.005-0.040-0 |
| Mutated R681 | 1.082 | 0.971 | 0.111 | 0.122 | 6 | R681-1.082-0.971-0.111-0.122-1 |
| Mutated G614 | 0.904 | 0.904 | 0.001 | 0.040 | 2 | G614-0.904-0.904-0.001-0.040-1 |
| DM G614-R681 |  |  |  |  |  |  |
| R681 | 1.023 | 0.975 | 0.047 | 0.066 | 6 | R681-1.023-0.975-0.047-0.066-1 |
| G614 | 0.901 | 0.901 | 0.001 | 0.041 | 2 | G614-0.901-0.901-0.001-0.041-1 |

The main observations of the data in **Table 1** are as follows:

1. AABP values provide the quantitative information on each AA position in the protein as the baseline comparisons to assess the mutation effect.
2. Significant differences in AABP values between site 681 and site 614 are noted. P681 has a much larger AABP than D614 due to their locations and interactions with other AAs.
3. Mutated R681 decreases the AABP by 1.082e^-^ - 1.117e^-^. = -0.035e^-^.
4. Mutated G614 decreases the AABP by 0.904e^-^ - 0.917e^-^ = -0.013e^-^.
5. The double mutation affects the changes in AABP for both sites: R681: 1.023e^-^ – 1.117e^-^ = -0.095e^-^. G614: 0.901 e^-^ – 0.917 e^-^ = -0.015e^-^. When single and double mutations are compared, the non-local AABP of R681 decreases by 0.047e^-^ – 0.111e^-^ = -0.064e^-^ in case of double mutation.
6. Please note that the contribution from the NL and HB part is a substantial portion of the total AABP.

To better explain the information present in **Table 1** regarding the nature of total AABP values and its components of nearest neighbor (NN) AABP and non-local AABP (NL) values, we show in **Figure 3** the sequence of AAs from E592 to S689. **Figure 3 (a)** in the ribbon form and **Figure 3 (b)** in the sequential form both showing the location of the main mutation sites P681 and D614 (pink circle), the NN AAs to these two mutations are in yellow circles and the non-local interacting AAs (green circle) connected by lines indicating that they are interacting.

**Figure 3.**
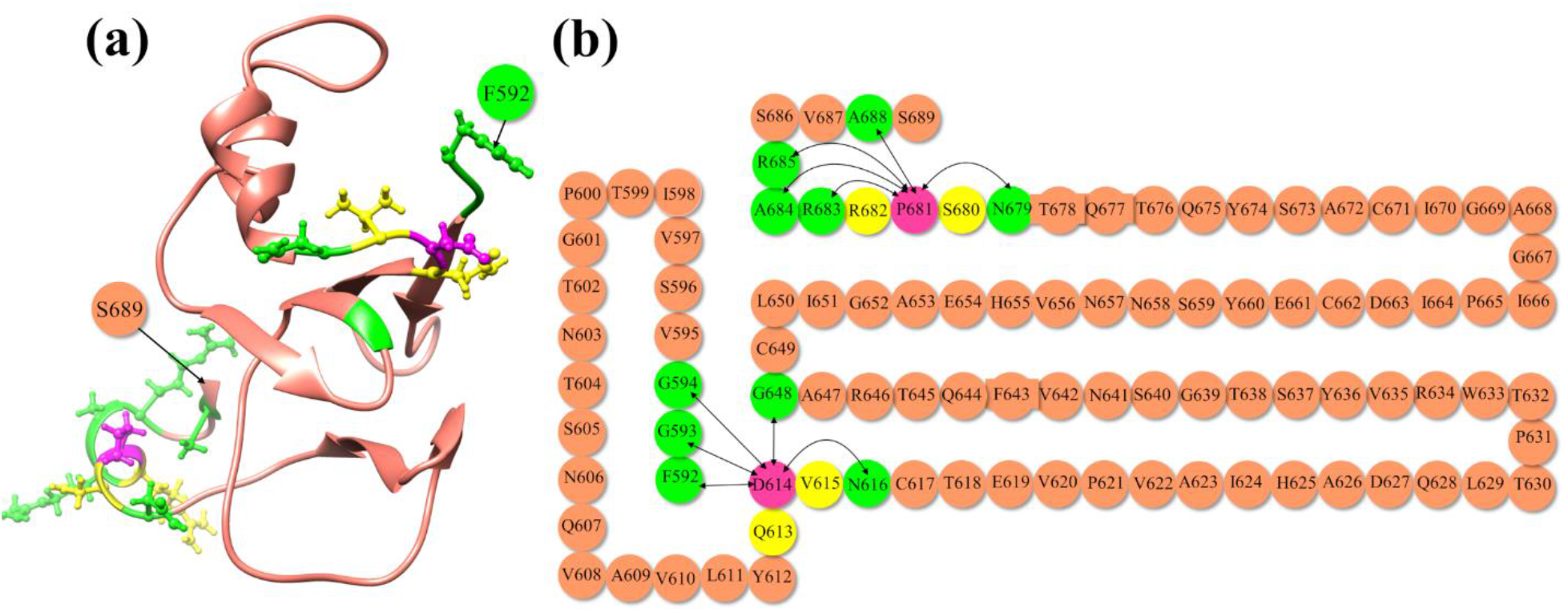
Interaction of D614 and P681 to their NN and NL AAs in WT SD2-FP model. **(a)** Ribbon structure from residue F592 to S689. **(b)** Sketch of AA sequence from F592 to S689 showing AABP interaction for D614 and P681 with joining lines. Both D614 and P681 are shown in pink color with its NN in yellow and NL interaction in green.

**Figure 4** shows more vividly the non-local AA-AA interactions of mutations in Delta variant. They are divided into six panels in two columns. (a) WT D614 and (b) WT P681; (c) mutated G614 and (d) mutated R681, and (e) double mutation G614-R681 G614 and (f) R681. In each case, the ball and stick sketch of all participating AAs are shown (red, O, grey, C, blue N, white, H). The focused AA is marked light pink. Its two NNs are marked light yellow, and its interacting NL AAs are marked light green. All interactions are marked by solid lines and the dashed lines show HBs. All these NL interactions with the bonds formed are listed in **Table S5, Table S6**, and **Table S7**. At the lower part of each figure, the three smaller figures show the same figure rotated for 90^°^, 180^°^, 270^°^ from left to right. These figures show some of the most detailed information on the AA-AA interaction at the atomic-scale summarized as follows.

**Figure 4.**
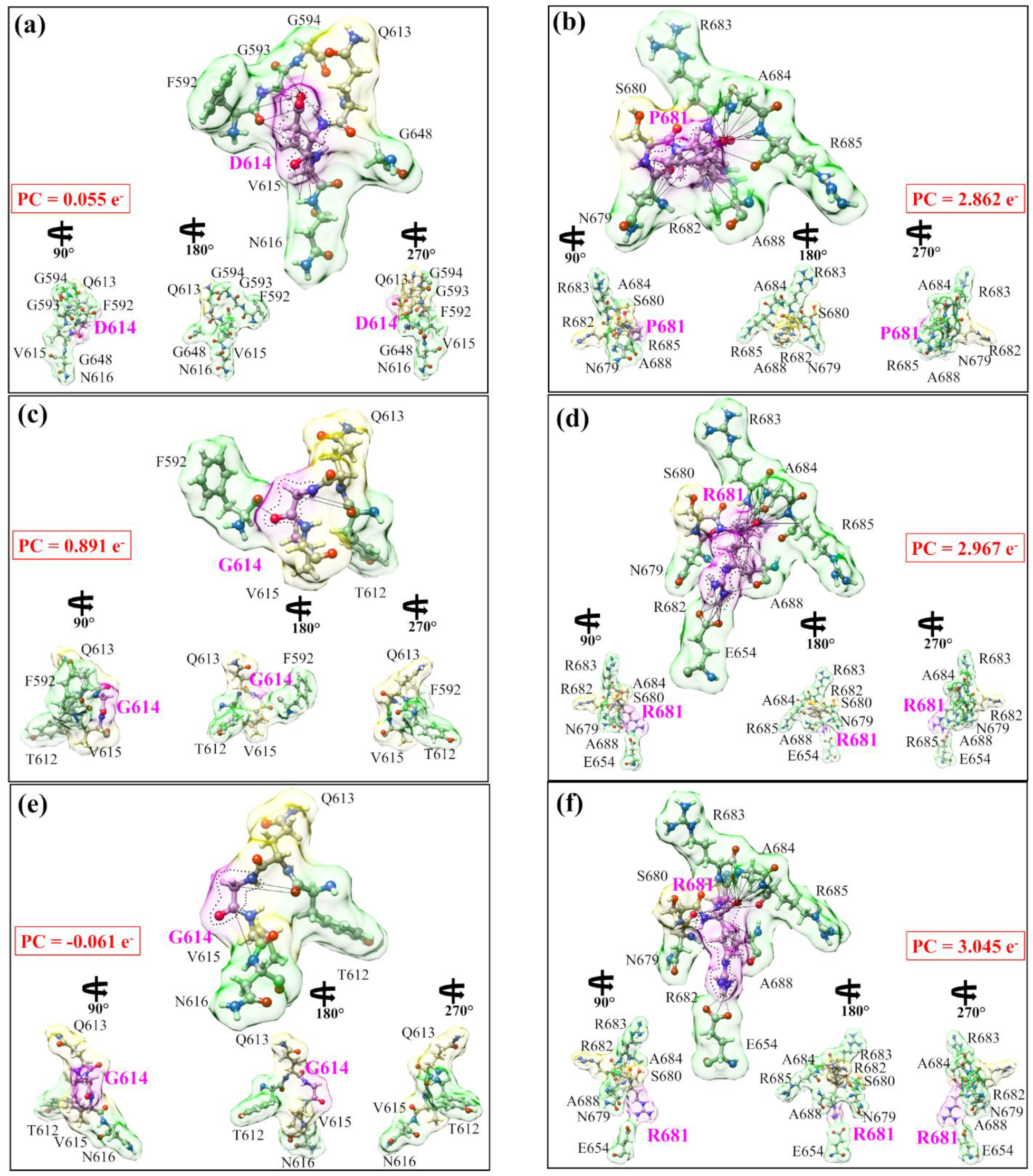
Details of the actual distribution of the three cases of interactions in the AABP calculation: (a) and (b) for the WT, (c) and (d) for the single mutation and (e) and (f) for the double mutation. The left panel in each case centered on 614^th^ site and the right panel centered on 681^st^ site both colored pink. The NN AAs are colored light yellow, and the NL AAs are colored light green. All the AAs involved in the interaction are marked. The lines show the NL bonded pairs. For each case, we show the NN AAs and the NL AAs that contribute to the AABP values. It also shows drastic difference between the mutation D614G and P681R in the shape, size and orientation and the total number of AAs involved in each case. The PC for each groups of AAs in the AABP calculation are listed in the red box showing large difference both in the sign and the magnitude.

(1) The WT D614 and WT P681 in (a) and (b) at the two different locations in the S-protein have very different features in size, shape and volume controlled by their interatomic interactions with the 2 NN and 5 NL AAs. (2) The mutated AAs G614 and R681 in (c) and (d) are drastically different from the WT case in (a) and (b). (3) The double mutation in (e) and (f) also show significant differences with WT and interacts with same number of NL AAs as in a single mutation.

These graphical illustrations demonstrate the complex structural features of mutations and interaction details never revealed before. In particular, the presence or the lack of HBs within the same AA or with other AAs have not been sufficiently elaborated in the current literature in biochemical interactions except in a few isolated cases. In the same figure, the partial charge on each group is shown in the red boxes. These PC values are obtained by summing PC values of individual AAs in each interacting group. In the WT, both groups involving D614 and P681 have positive PC. After mutation, G614 and its interacting group has its PC significantly increase whereas R681 and its interacting group has positive PC only slightly increased. More surprisingly, in the case of double mutation, G614 with its group have its PC slightly negative and P681 with its group have its positive PC continue to increase. This is another solid evidence for the strong effect of mutation on different AAs that has never been discussed or revealed before. Similarly, the detailed interaction of 950^th^ site in two HR1-CH models is shown in **Figure S7** with their AABP are listed in **Table S8**, their NL bonding listed in **Table S9**, and their results are discussed in **section S2** in **SI**.

## 5. DISCUSSIONS

### 5.1 The origins of mutation

Viruses can undergo frequent genetic mutations, including point mutations (the source of genetic variation) and recombination ^58^. Mutations are nucleotide changes that result in AA sequence changes implying new phenotype variants, whereas recombination allows these variants to move across genomes to produce new haplotypes. Recombination occurs when viruses containing variants with different mutations infect the same host cell and exchange the genetic segments ^58^. The fate of these genetic changes will be ultimately determined by natural selection and genetic drift, so it is very difficult to forecast when a viral mutation will become globally dominant. Although coronaviruses have an exonuclease enzyme that reduces their replication error rates, they accumulate mutations and generate more diversity via the recombination ^57, 59, 60^. SARS-CoV-2 itself is most likely the result of a recombination between different SARS-related coronaviruses ^61^, with different subsequent mutations affecting its many biological and biomedical properties such as pathogenicity, infectivity, transmissibility, and/or antigenicity even though they tend to be either deleterious and quickly purged, or relatively neutral ^62^.

One of the most urgent tasks in the virus research is the origin of virus mutations and how to predict new variants even before they would occur. We assert that the first task must be related to the interaction between AAs at the atomic level that result in the structural modification in the S-protein due to mutated AAs. It must involve the interaction with non-local AAs in addition to the NN AAs. The contribution to AABP from hydrogen bonding is critical and the role of HB has been recognized by all researchers but seldom explored in detail. In a much broader sense, the origin of mutation is not limited just to SARS-CoV-2 research *per se* but is related to broader themes of evolutionary biology such as the origin of species ^63^. This accentuates the importance of a fundamental understanding of biomolecular interaction.

The data in **Table 1** reveals that D614G and P681R mutations have lower AABP values than unmutated WT case. Our results make sense for the following reasons. First, the substitution of D614 with G results in losing the sidechain, leading to the elimination of many intramolecular interactions in the same protomer as shown in **Figure 4**. More specifically, our result elucidates that this mutation disrupts the non-local network interactions (**Table 1**). This could have large structure consequences in other domains of S-protein such as in promoting the up conformation of S-protein or enhancing cleavage site as reported before ^29, 30^. This enhancement in the cleavage site could be due to an increase in flexibility of the mutated 614 and 681 sites. In addition to these intra-protomer interactions, it has been structurally demonstrated that the D614G mutation destroys an inter-protomer hydrogen bond between D614 (chain A) and T859 (chain B) ^64^. However, the SD2-FP model alone is insufficient to assess these significant conformational changes. It is necessary to include all atoms of the S protein trimer, which is currently impossible to perform in a single *ab initio* calculation. Second, our decomposition of the total AABP into NN and NL AABPs reveals that the R681 increases the NL AABP by forming new HBs and decreases the NN AABP as compared to P681 (**Table 1** and **Figure 4**). Importantly, P681R mutation *reduces the local rigidity as evidenced by the NN AABP values* (**Table 1**). Additionally, the proline is well-known as the most rigid AA, and when it is mutated to arginine at position 681, it loses its rigidity. Furthermore, the positive charge associated with R681 appears to alter virus tropism via enhancing S1/S2 cleavage, as previously demonstrated in human airway epithelial cells ^28^. This coincides with our conclusion that mutation decreases AABP but also increases the flexibility of AAs.

Such understanding of the molecular and atomic origins of the individual mutations or their combinations in SARS-CoV-2 can provide deep information to prepare and prevent future outbreaks such as those reported for AY.4.2. or the just emerging “Omicron” VOC. It can also play an important role in guiding the development of new drugs. It would be also desirable to have a quantitative scale from 1 (insignificant) to 10 (most dangerous) to quantify the nature of mutations by linking the mutation to specific clinical data or research using other methods, experimental or computational.

### 5.2 Extension to machine learning (ML)

Over the last decade or so machine learning (ML) has become a very powerful tool that is being applied to many different areas including image, speech, text and facial recognition, autonomous vehicles, medical image classification, instruction detection, finances, drones, and national defense, etc. ^65, 66^. In the present work, the word ML is strictly used only for the calculated data of AABP between AAs in the S-protein to predict potential unknown mutations.

One of the major challenges we face in applying ML techniques to our problem is the size of the dataset. This is because each data instance of a Delta variant model is computationally expensive to generate. When dealing with small datasets, deep learning techniques (based on artificial neural networks) will tend to under-perform. Hence, conventional ML techniques should be preferred. Our first step is to prepare the data by constructing feature vectors for different Delta variant models. We can represent each model as a 5-dimensional feature vector: **TAABP** (Total AABP), **NN** (NN AABP), **NL** (Non-local AABP), **HB** (AABP from HB), and **NNL** (No. of NN AAs). Each feature is a continuous variable. The target vector is denoted by a binary variable **MT** to represent weather a Delta variant model is unmutated (0) or mutated (1). **Table 2** shows how the data can be represented for the Delta variant models shown in **Tables 1** for SD1-FP and **Table S8** for HR1-CH. Several extensions can be made to the target vector depending on the prediction task. Suppose we wish to predict the site of the Delta variant model (e.g., 614, 681), a categorical variable can be introduced as the target vector. To predict different kinds of mutations (e.g., single, double), another categorical variable can be introduced in the target vector.

**Table 2.**
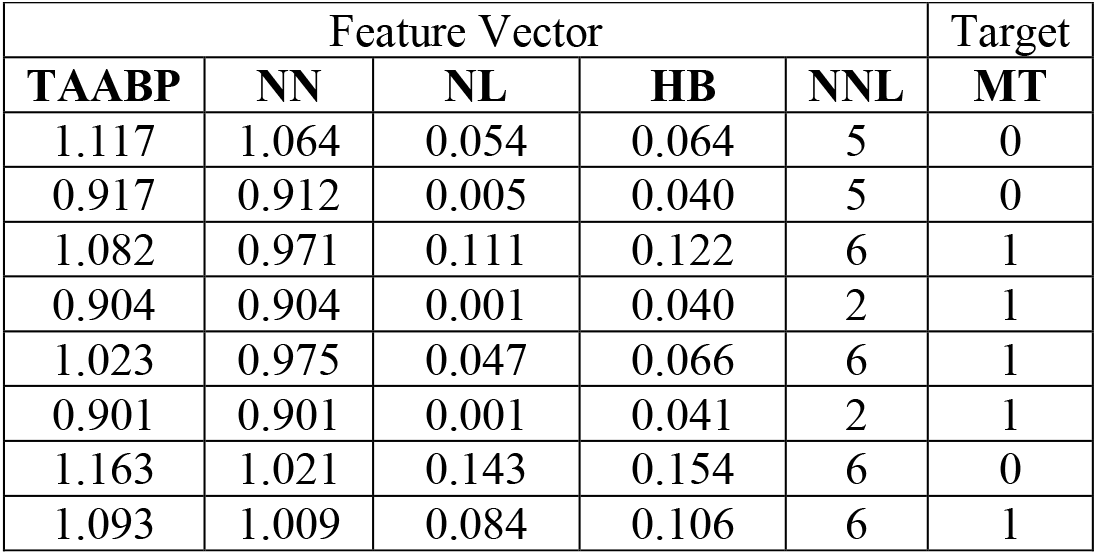
Data representation for ML.

Suppose we wish to predict whether a new Delta variant model is mutated or not using the feature and target vectors in **Table 2**, a binary classifier can be learned on the data. Classifiers can be built using different techniques such as (a) Logistic Regression classifier ^67^, (b) Gaussian Naïve Bayes classifier ^67^, (c) Support Vector Machine (SVM) classifier ^68^, (d) Decision Tree classifier ^69^, (e) Random Forest classifier ^70^, and (f) the extreme Gradient Boosting (XGBoost) classifier ^71^. Hyperparameter tuning is necessary for the different classifiers to achieve the best accuracy. Feature importance is another task that can be valuable to researchers. For instance, the XGBoost classifier trained on data in **Table 2** showed that **TAABP** and **HB** were the most important features to predict a mutation. XGBoost uses the notion of gain, which is the relative contribution of a feature to the model, to compute feature importance. The above ML techniques assume that the data instances are independently and identically distributed (i.i.d). However, if this assumption is not appropriate, then ML techniques such as Markov logic networks ^72^ should be considered. We can also learn causal relationships among the features using a probabilistic graphical model such as a Bayesian network ^73^. By better understanding the cause-effect relationships among the features, better prediction models can be developed. A Bayesian network can compactly encode the joint distribution of a set of random variables as the product of the conditional probability of the nodes given its parents in the network. Using a Bayesian network, we can also simulate new data that follows the distribution learned by the network. **Table 3** shows a set of 4 samples obtained by simulating random samples from a Bayesian network. (This network was learned using the Max-Min Hill Climbing algorithm ^74^ over the data shown in **Table 2**.) This algorithm first learns the skeleton of a Bayesian network. It then uses greedy hill-climbing search to orient the edges using a Bayesian scoring function ^74^.)

**Table 3.**
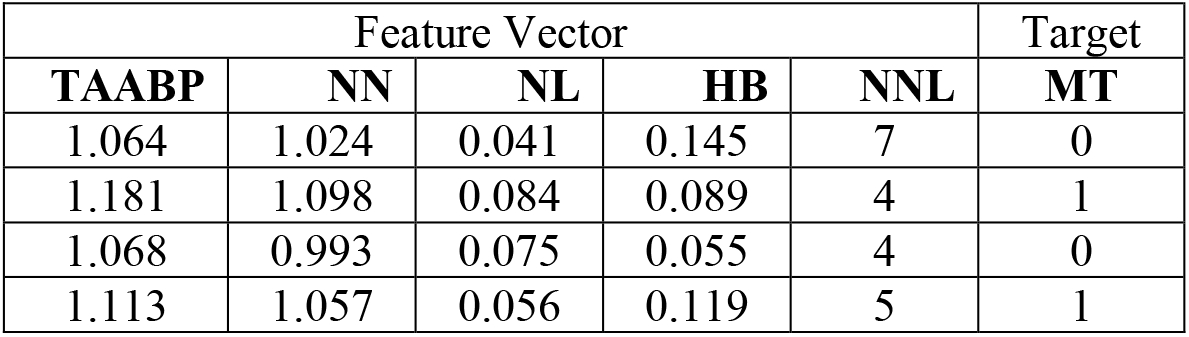
Data obtained by simulating random samples from a Bayesian network

Figure 5 shows the general steps involved in applying supervised ML to our Delta variant data. In summary, ML can offer unprecedented opportunities to predict mutations in Delta variant.

**Figure 5.**
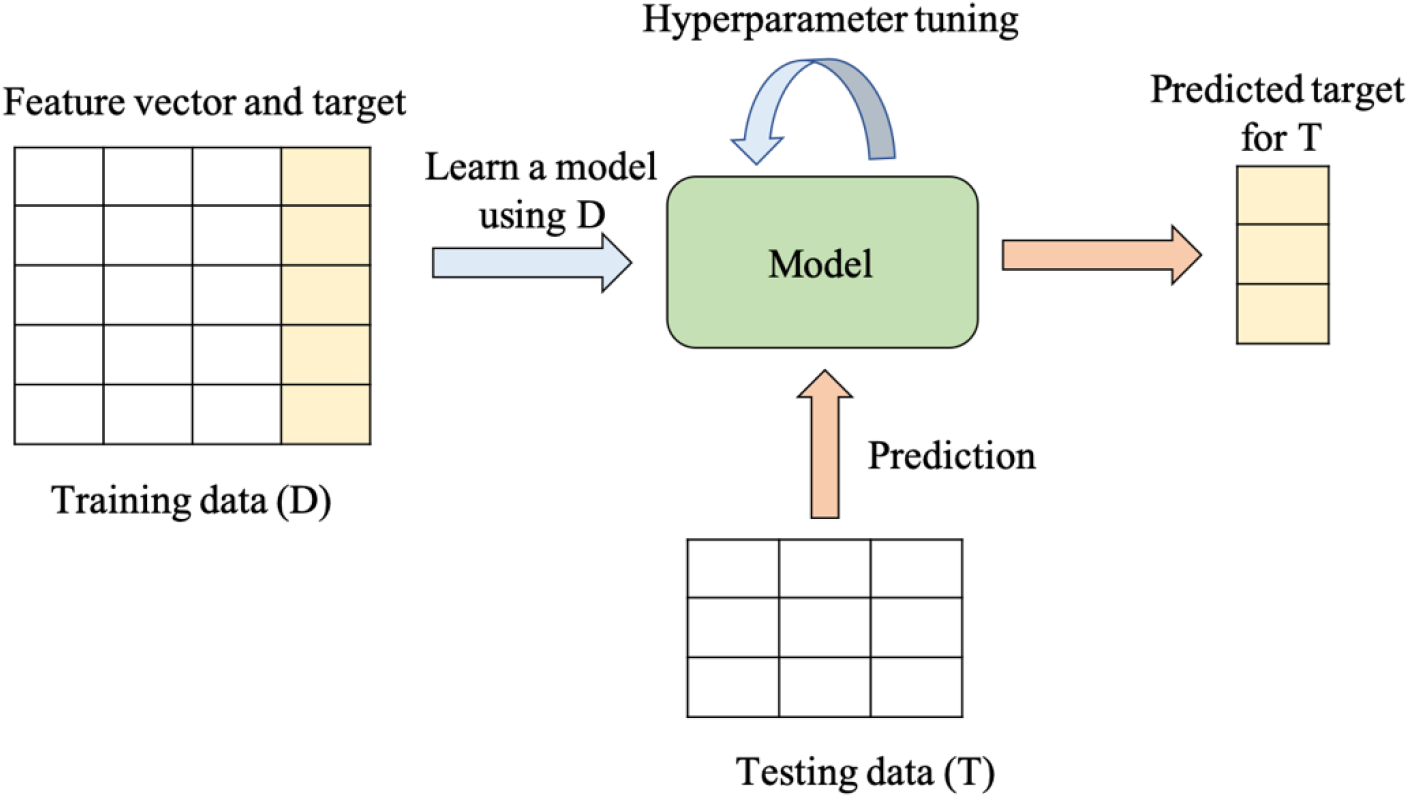
The general workflow of how supervised ML works is shown. Here D and T represents training data and testing data respectively.

### 5.3 Looking forward

Looking forward, we would like to speculate where our methodology could lead in the future. Obviously, it can be extended to other mutations in S-protein such as RBD and NTD and their interfaces with ACE2 or maybe even those newly emerged mutations such as AY.4.2 ^17^ and B.1.1.529 ^75^. The obvious goal is to increase the mutation data points so a reasonable size of the AABP database can be used in the ML as discussed in the previous section. It is totally possible to calculate other potential mutations for more data points in different domains based on the insights obtained with the calculations on known mutations. The list is endless, restricted by the large computational resources it demands. Fortunately, the emergence of the next generation of exa-scale supercomputer is already available ^76^ to meet these challenges.

However, even more importantly, we posit that our methodology could be extremely valuable for the VOC that appear to be characterized by an unprecedented number of concurrent mutations - as appears to be the case in the latest Omicron VOC ^75^ - with more than 30 mutations per S-protein. Clearly in such cases there must be collective effects tying together several mutations, possibly also enhancing the susceptibility of different AAs to mutations. The mutation quantifier based on AABP, as introduced in this paper, which by its very nature already embodies the collective effects of mutations, is clearly a good candidate for such an analysis. In view of this striking development of SARS-CoV2 mutational capacity, our research seems to represent a well-placed origin for further elaboration not only of the effects of single mutations, but maybe even more importantly - their *interaction and synergy*.

One of the fundamental questions in biology is to ask if mutations are random or if there are specific reasons for each mutation. We may be able to shed some light also on this issue by accumulating a large database for known mutations and applying the ML protocol to see if any successful prediction can be verified with a global database or clinical data. So far, we have 6 data points plus another 4 from RBD, or 10 data points altogether.

The current calculation and analysis are restricted to Chain A of the S-protein of SARS-CoV-2. Extension to cases with chains A, B, and C in up and down conformations would be highly desirable and important. It is also opportune to start looking at mutations in RBD of S-protein such as those involved in binding to the ACE2 receptor to provide a deeper understanding of how the virus can mutate and overcome the human defense mechanisms of immune response, intimately related to vaccination. Most neutralizing monoclonal antibodies (mAbs) target either the RBD or the NTD of S-protein, and mutations in these domains have the greatest impact on neutralization.

The broader implications of the present work would be also in protein-protein interactions where one could define similar *ab initio* interaction quantifiers based on single-point calculations as applied to protein-protein network, protein-protein mapping, application to cancer research etc. This broader phenomenology also revolves around mutations (replacing AA at a specific site, some AAs are more important than others) but on a much larger scale and with much more detailed interactions. While so far, such studies were essentially based on experimental or clinical observations, the lack a firm fundamental theoretical basis is noticeable.

## ASSOCIATED CONTENT

## SUPPORTING INFORMATION

Additional descriptions, figures and Tables are provided in the Supporting Information.

## AUTHOR INFORMATION

### Author Contributions

WC and PA conceived the project. WC and PA performed the calculations. PA and BJ made most of the figures. WC, PA, BJ drafted the paper with inputs from RP and PR, RG provided the computational resources. All authors participated in the discussion and interpretation of the results. All authors edited and proofread the final manuscript.

## ACKNOWLEDGMENTS

This research used the resources of the National Energy Research Scientific Computing Center supported by DOE under Contract No. DE-AC03-76SF00098 and the Research Computing Support Services (RCSS) of the University of Missouri System. We thank Dr. Richard Gerber, Senior Science Advisor and HPC Department Head for special allocations. This project is funded partly by the National Science Foundation of USA: RAPID DMR/CMMT-2028803. P. Rao was partly funded by the National Science Foundation of USA: RAPID CISE/CNS-2034247.RP acknowledges funding from the Key project #12034019 of the National Natural Science Foundation of China.

