## Supporting information for "Delta variant with P681R critical mutation revealed by ultra-large atomic-scale *ab initio* simulation: Implications for the fundamentals of biomolecular interactions"

### S1. COMPUTATIONAL METHODS.

**S1.1 Vienna *ab initio* simulation package (VASP).** The initial structure for the WT and mutation models mentioned in Section 2.1 are then optimized by using the *Vienna ab initio simulation package* (VASP), which is known for its efficiency in structural relaxation based on the *density functional theory* (DFT). We use the projector augmented wave (PAW) method with Perdew-Burke-Ernzerhof (PBE) exchange correlation functional [1] within the generalized gradient approximation (GGA). Our experience and detailed tests suggest the use of the following input parameters for biomolecular systems as more than adequate: energy cut-off 500 eV; electronic convergence of $10^{-3}$ eV; force convergence criteria for ionic steps at $-10^{-2}$ eV/Å; and single k-point sampling. Details of the test is shown in **Table S2** and discussed in the **Results section 4**. All structure optimizations using VASP have been carried out at the National Energy Research Scientific Computing (NERSC) facility using Cori at the Lawrence Berkeley Laboratory or at the Research Computing Support Services (RCSS) of the University of Missouri System. The VASP-optimized structure is used as the input for the electronic structure and interatomic bonding calculations described below.

**S1.2 Orthogonalized linear combination of atomic orbitals (OLCAO) method.** For the electronic structure and interatomic interactions in biomolecules, we use a very different DFT method, the all-electron *orthogonalized linear combination of atomic orbitals* (OLCAO) method developed in-house. The efficacy of using the combination of these two different DFT codes is well documented [2-15]. The key feature of the OLCAO method is the provision for the effective charge ($Q^*$) on each atom and the *bond order* (BO) values $\rho_{\alpha\beta}$ between any pairs of atoms. They are obtained from the *ab initio* wave functions with atomic basis expansion calculated quantum mechanically

$$Q_\alpha^* = \sum_i \sum_{m,occ} \sum_{j,\beta} C_{i\alpha}^{*m} C_{j\beta}^{m} S_{i\alpha,j\beta} \quad (1)$$

$$\rho_{\alpha\beta} = \sum_{m,occ} \sum_{i,j} C_{i\alpha}^{*m} C_{j\beta}^{m} S_{i\alpha,j\beta} \quad (2)$$

In the above equations, $S_{i\alpha,j\beta}$ are the overlap integrals between the $i^{th}$ orbital in $\alpha^{th}$ atom and the $j^{th}$ orbital in the $\beta^{th}$ atom. $C_{j\beta}^{m}$ are the eigenvector coefficients of the $m^{th}$occupied molecular orbital. The *partial charge* (PC) or ($\Delta Q_{\alpha} = Q_{\alpha}^{0} \ - \ Q_{\alpha}^{*}$ ) is the deviation of the effective charge $Q_{\alpha}^{*}$ from the neutral atomic charge $Q_{\alpha}^{0}$ on the same atom $\alpha$. The BO quantifies the strength of the bond between two atoms and usually scales with the *bond length* (BL) but is also influenced by the surrounding atoms. The calculations of PC and BO is based on the Mulliken scheme [16, 17], hence are basis-dependent, so that comparisons of BO values using different basis sets or different DFT methods should be treated with caution. We use a one-point calculation to characterize the bonding rather than traditional energy (or enthalpy) difference calculation (two-point calculation) which requires reference energy for comparison and depends on the method used. The sum of all BO values within a single structure unit gives the *total bond order* (TBO). This is an innovative aspect of our method in the study of complex biomolecular materials capable to yield a large amount of crucially important data.

### S2. ADDITIONAL SUPPORTING INFORMATION.

(a) We have added the AABP data for a single mutation D950N in domain HR1-CH and listed in **Table S8**. The AABP data essentially confirm the observations listed in the previous **subsection 4.3** that mutation reduces the AABP values and weakens the bonding between amino acids. Both WT and mutated cases have the same number of NL interactions (see **Figure S7**). However, the bonding in each case is different as shown in **Table S9.** Comparing **Figure S7** for mutation D950N in HR1-CH with the main **Figure 4** for mutations D614G and P681R in SD2-FP, we can clearly see that (1) The overall shape and size of WT D950 have very small difference with the mutated N950; (2) These results are very different from mutations D614G and P681R in SD2-FP in **Figure 4**; (3) The PC of WT D950 and its interacting group (+0.209e$^-$) is significantly increased in case of mutated N950 and its interacting group(+1.146e$^-$). These observations again support our assertion that the effect on mutation critically depends on the location of the targeted residue and its interactions with its neighboring AAs, NN and NL ones, via the detailed atomic scale calculation of AABP values.

(b) We have PC for each AAs of WT HR1-CH listed in **Table S4** and bar graph in **Figure S6**. **Figure S6** shows PC of AA D950 changes from highly negative -0.813 e$^-$ to 0.073 e$^-$ when mutates to N950.

### Supporting Figures:

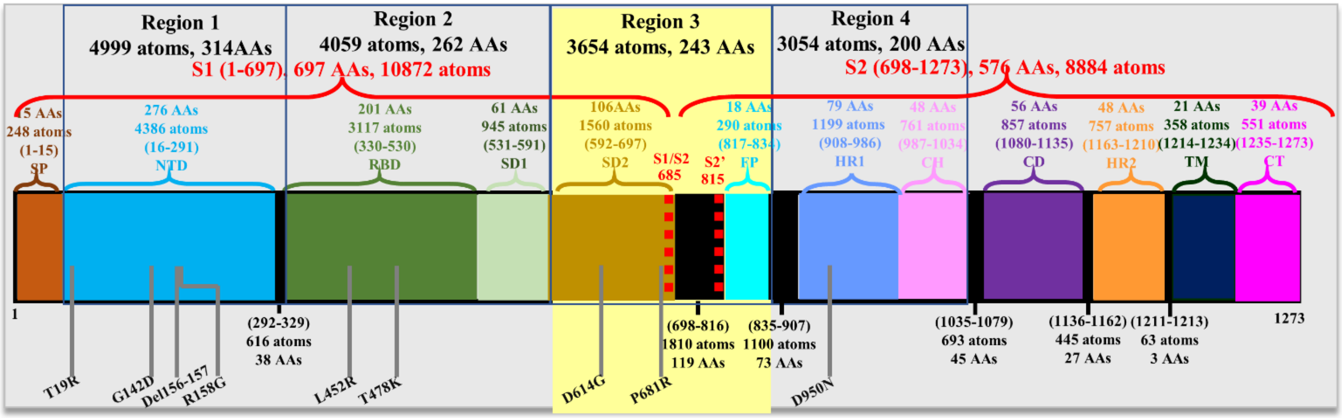


**Figure S1.** Detailed schematic of S-protein in SARS-CoV-2 colored by domains: SP, NTD, RBD, SD2, furin cleavage site (S1/S2), FP, HR1, CH, CD, HR2, TM, and CT. The delta variant mutation sites are marked by gray solid line in the bottom. In the present work, our calculations have been mainly carried out on region 3 of SD2 and FP domains.

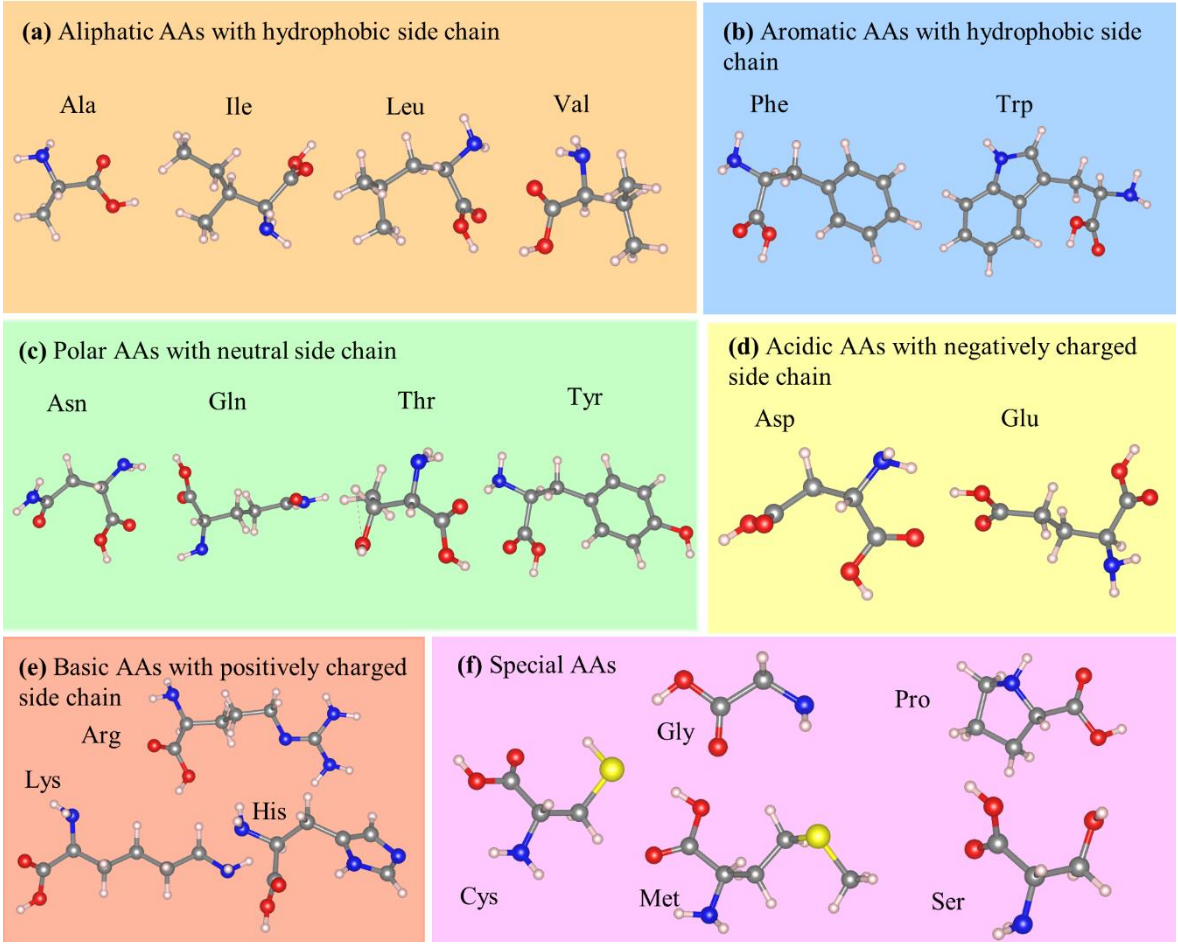


**Figure S2.** The 20 amino acids in **Table S1** are divided into 6 functional groups based on the nature of their sidechains: (a) and (b) for hydrophobic AAs, (c) for neutral polar AAs, (d) and (e) for charged polar AAs, and (f) for unique AAs.

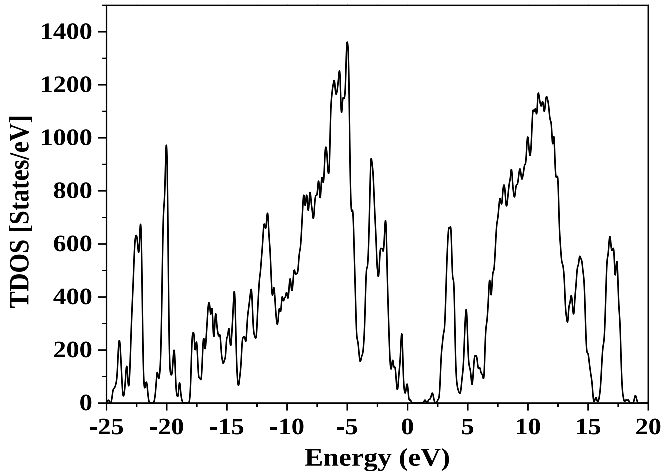


**Figure S3.** Calculated total density of stated (TDOS) for WT SD2-FP model.

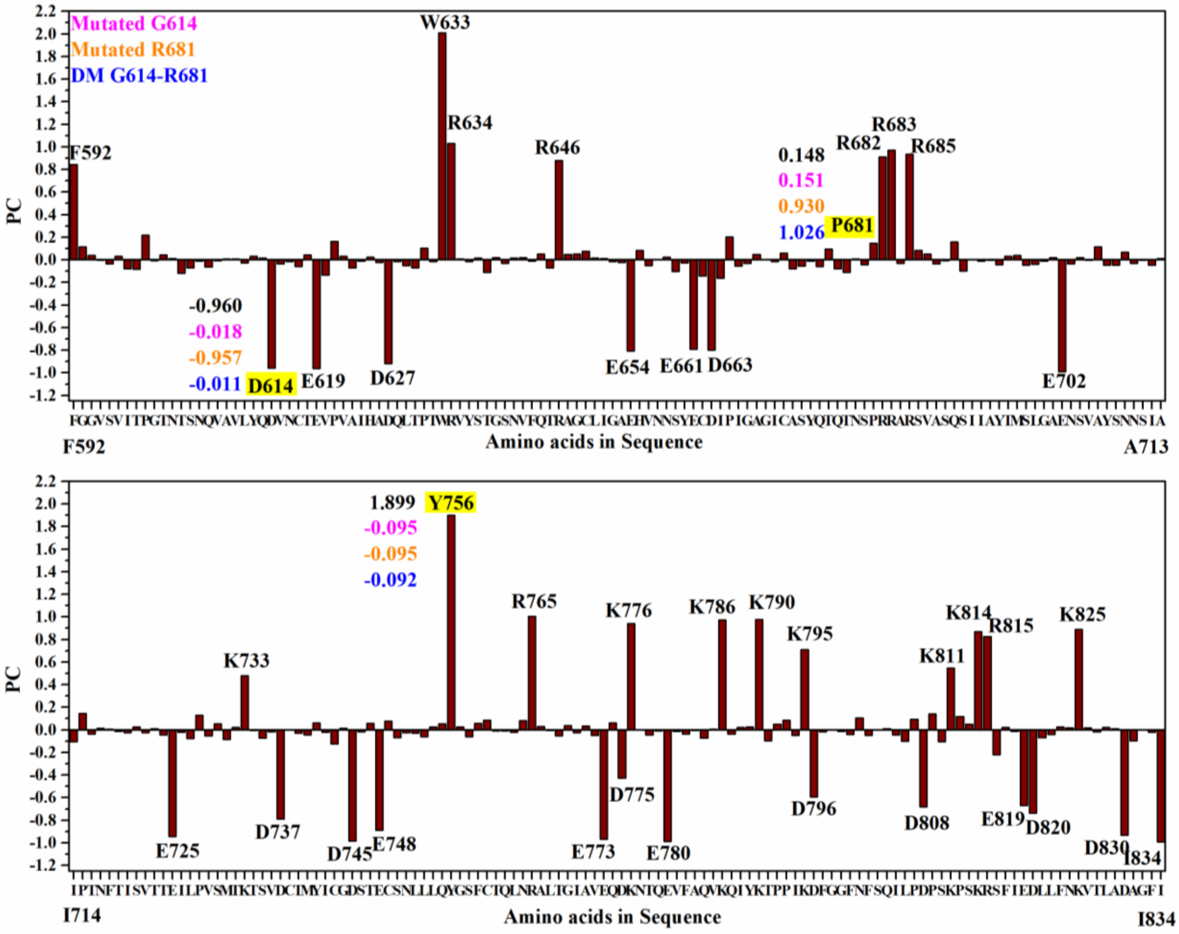


**Figure S4.** Bar graph with PC distribution for WT SD2-FP. PC values are marked for the two mutation sites 614 and 681 showing values in different color for different mutation cases.

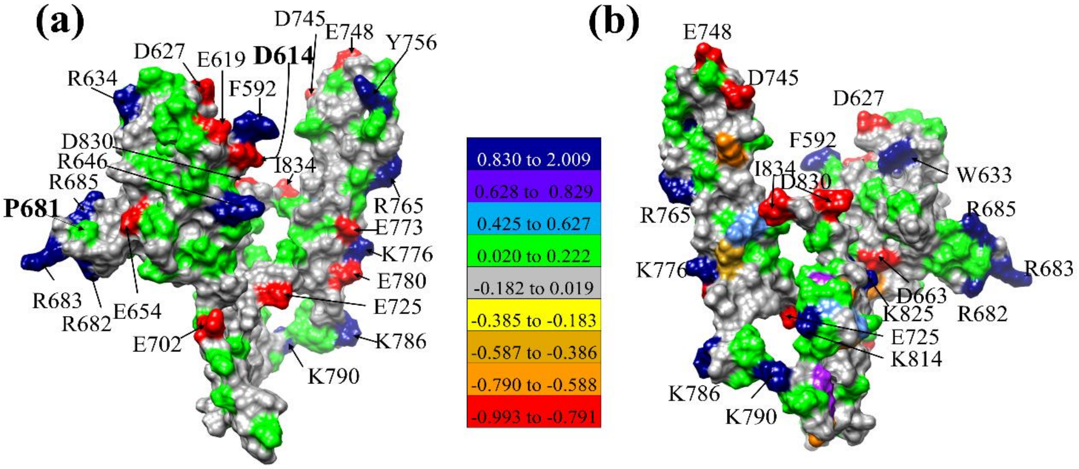


**Figure S5. (a)** Comparison of PC on the solvent excluded surface between P681 and D614**. (b)** 180° orientation of **(a)**.

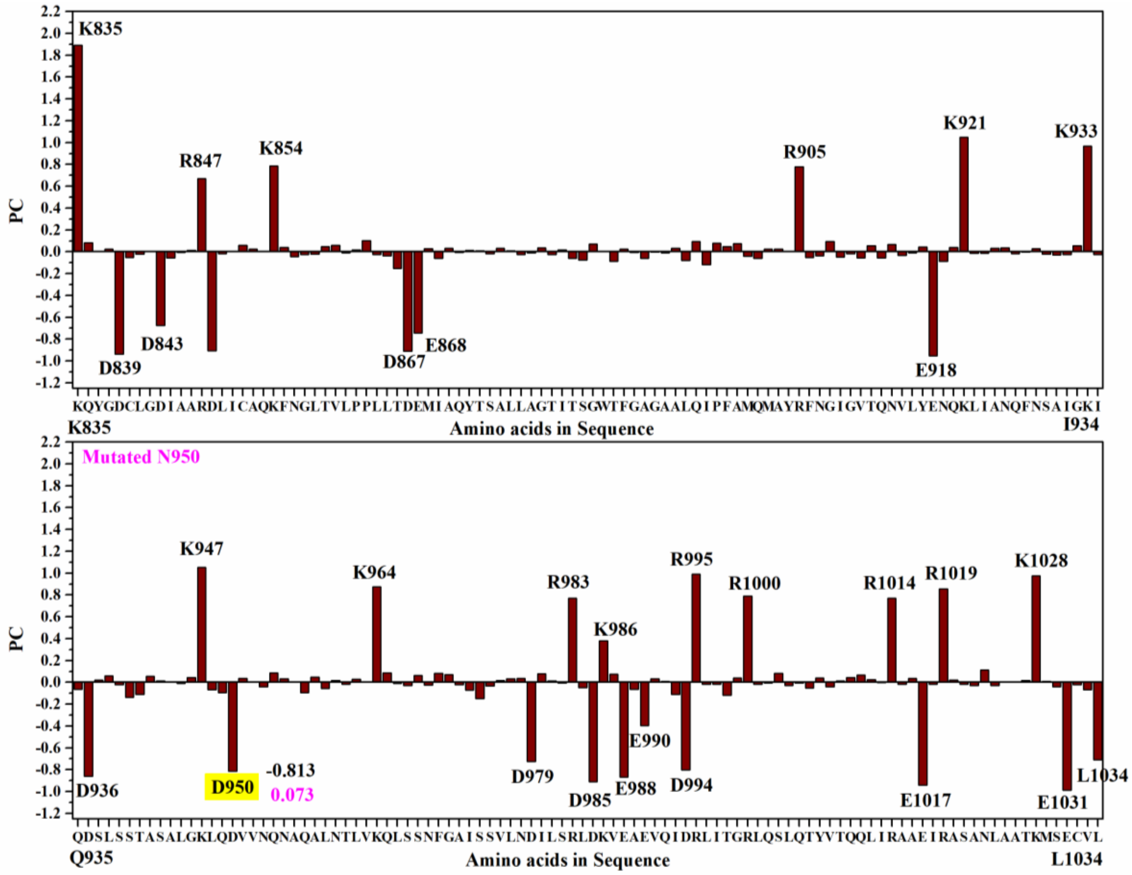


**Figure S6.** Bar graph with PC distribution for WT HR1-CH. PC values are marked for the two mutation sites 950 showing values in different color for D950N mutation.

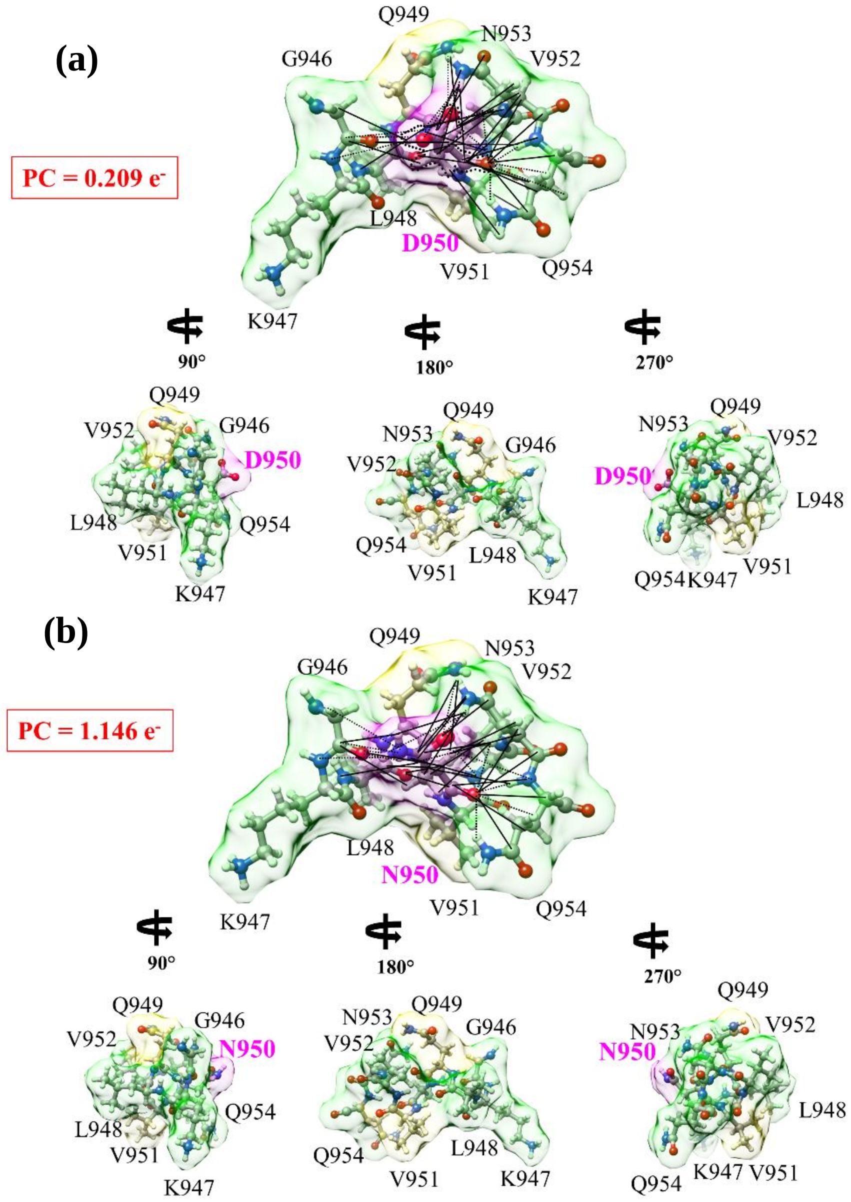


**Figure S7.** Interactions of 950$^{th}$ site in two HR1-CH models in Delta variant: (a) WT D950 and (b) mutated N950. In each case, the ball and stick sketch of all participating AAs are shown (red, O, grey, C, blue N, white, H). The focused 950$^{th}$ AA is marked light pink. Its two NNs are marked light yellow, and its interacting NL AAs are marked light green. All NL interactions are marked by solid lines and dashed lines show HBs. All these NL interactions with the bonds formed are shown in **Table S9.** At the lower part of each of the figure, three smaller figures show the same figure rotated for 90˚, 180˚, 270˚ form left to right. These figures show some of the most detailed information on the AA-AA interaction at atomic scale. In the same figure, the partial charge on each group is shown in the red boxes. These PC values are obtained summing PC values of individual AAs in each interacting group. The mutated case has distinctly higher positive PC.

**Supporting Tables:**

**Table S1.** 20 canonical amino acids in alphabetical order. The last column shows their functional group.

| No. | Amino acid | 3 letter code | 1 letter code | TBO | No of Atoms | TBO /Atom | No. of bonds | No. of HB | Group |
|---|---|---|---|---|---|---|---|---|---|
| 1 | **Alanine** | **Ala** | **A** | 4.380 | 13 | 0.337 | 25 | 5 | a |
| 2 | **Arginine** | **Arg** | **R** | 9.434 | 26 | 0.363 | 52 | 12 | e |
| 3 | **Asparagine** | **Asn** | **N** | 6.016 | 17 | 0.354 | 30 | 7 | c |
| 4 | **Asparatic** | **Asp** | **D** | 5.456 | 16 | 0.341 | 33 | 7 | d |
| 5 | **Cystein** | **Cys** | **C** | 4.512 | 14 | 0.322 | 23 | 3 | f |
| 6 | **Glutamine** | **Gln** | **Q** | 7.224 | 20 | 0.361 | 38 | 9 | c |
| 7 | **Glutamic acid** | **Glu** | **E** | 6.664 | 19 | 0.351 | 37 | 7 | d |
| 8 | **Glycine** | **Gly** | **G** | 3.198 | 10 | 0.320 | 16 | 4 | f |
| 9 | **Histidine** | **His** | **H** | 8.164 | 20 | 0.408 | 41 | 7 | e |
| 10 | **Isoleucin** | **Ile** | **I** | 8.033 | 22 | 0.365 | 43 | 6 | a |
| 11 | **Leucin** | **Leu** | **L** | 8.032 | 22 | 0.365 | 40 | 4 | a |
| 12 | **Lysine** | **Lys** | **K** | 8.771 | 24 | 0.365 | 44 | 7 | e |
| 13 | **Methonine** | **Met** | **M** | 6.879 | 20 | 0.344 | 35 | 2 | f |
| 14 | **Phenylalanine** | **Phe** | **F** | 10.119 | 23 | 0.440 | 47 | 3 | b |
| 15 | **Proline** | **Pro** | **P** | 6.488 | 17 | 0.382 | 42 | 6 | f |
| 16 | **Serine** | **Ser** | **S** | 4.481 | 14 | 0.320 | 27 | 4 | f |
| 17 | **Threonine** | **Thr** | **T** | 5.642 | 17 | 0.332 | 31 | 5 | c |
| 18 | **Tryptophan** | **Trp** | **W** | 12.436 | 27 | 0.461 | 53 | 2 | b |
| 19 | **Tyrosine** | **Tyr** | **Y** | 10.248 | 24 | 0.427 | 47 | 3 | c |
| 20 | **Valine** | **Val** | **V** | 6.758 | 19 | 0.356 | 36 | 4 | a |

**Table S2**. Four SD2-FP models ((a) to (d)) and two HR1-CH models ((e) and (f)) with the number of atoms, total energy and time used in Cori.

| Models | Atoms | Total Energy (eV) | Convergence | Time in Cori (node hour) |
|---|---|---|---|---|
| (a) WT P681 & D614 | 3654 | -22293.12593 | -0.0479275 | 12x48x64= 36864 |
| (b) Mutated R681 | 3664 | -22350.51693 | -0.0339289 | 13x48x64= 39936 |
| (c) Mutated G614 | 3649 | -22259.50298 | -0.0339803 | 13x48x64= 39936 |
| (d) DM G614-R681 | 3659 | -22316.36542 | -0.0324175 | 12x48x64=36864 |
| (e) WT D950 | 3054 | -18528.45614 | -0.00613611 | 6x48x64= 18432 |
| (f) Mutated N950 | 3056 | -18538.10922 | -0.00622162 | 5x48x64=15360 |

**Table S3:** List of PC value for each amino acids with their sequence number for WT SD2-FP.

| AA Seq No | PC | AA Seq No | PC | AA Seq No | PC | AA Seq No | PC | AA Seq No | PC |
|---|---|---|---|---|---|---|---|---|---|
| F592 | 0.843 | N641 | 0.017 | Q690 | 0.158 | T739 | -0.030 | I788 | 0.021 |
| G593 | 0.116 | V642 | 0.021 | S691 | -0.099 | M740 | -0.046 | Y789 | 0.027 |
| G594 | 0.040 | F643 | -0.013 | I692 | 0.005 | Y741 | 0.061 | K790 | 0.977 |
| V595 | -0.003 | Q644 | 0.051 | I693 | -0.013 | I742 | -0.022 | T791 | -0.098 |
| S596 | -0.033 | T645 | -0.070 | A694 | -0.003 | C743 | -0.123 | P792 | 0.050 |
| V597 | 0.032 | R646 | 0.878 | Y695 | -0.042 | G744 | 0.013 | P793 | 0.085 |
| I598 | -0.080 | A647 | 0.047 | T696 | 0.034 | D745 | -0.986 | I794 | -0.050 |
| T599 | -0.084 | G648 | 0.051 | M697 | 0.042 | S746 | -0.018 | K795 | 0.711 |
| P600 | 0.218 | C649 | 0.074 | S698 | -0.048 | T747 | 0.060 | D796 | -0.595 |
| G601 | -0.008 | L650 | 0.016 | L699 | -0.040 | E748 | -0.889 | F797 | -0.018 |
| T602 | 0.046 | I651 | 0.013 | G700 | -0.013 | C749 | 0.079 | G798 | -0.003 |
| N603 | 0.011 | G652 | -0.017 | A701 | 0.021 | S750 | -0.069 | G799 | -0.015 |
| T604 | -0.118 | A653 | -0.024 | E702 | -0.991 | N751 | -0.025 | F800 | -0.041 |
| S605 | -0.070 | E654 | -0.808 | N703 | -0.035 | L752 | -0.030 | N801 | 0.104 |
| N606 | -0.013 | H655 | 0.085 | S704 | 0.020 | L753 | -0.061 | F802 | -0.050 |
| Q607 | -0.065 | V656 | -0.051 | V705 | -0.004 | L754 | 0.027 | S803 | -0.001 |
| V608 | -0.009 | N657 | 0.000 | A706 | 0.114 | Q755 | 0.053 | Q804 | 0.012 |
| A609 | 0.010 | N658 | 0.022 | Y707 | -0.046 | Y756 | 1.899 | I805 | -0.046 |
| V610 | 0.009 | S659 | -0.103 | S708 | -0.046 | G757 | 0.026 | L806 | -0.099 |
| L611 | -0.029 | Y660 | -0.029 | N709 | 0.066 | S758 | -0.061 | P807 | 0.092 |
| Y612 | 0.032 | E661 | -0.790 | N710 | -0.033 | F759 | 0.057 | D808 | -0.680 |
| Q613 | 0.016 | C662 | -0.144 | S711 | -0.005 | C760 | 0.085 | P809 | 0.143 |
| D614 | -0.960 | D663 | -0.799 | I712 | -0.047 | T761 | -0.009 | S810 | -0.104 |
| V615 | -0.036 | I664 | -0.162 | A713 | 0.013 | Q762 | -0.008 | K811 | 0.549 |
| N616 | -0.017 | P665 | 0.204 | I714 | -0.104 | L763 | -0.021 | P812 | 0.119 |
| C617 | -0.058 | I666 | -0.055 | P715 | 0.145 | N764 | 0.084 | S813 | 0.049 |
| T618 | 0.045 | G667 | -0.031 | T716 | -0.039 | R765 | 1.006 | K814 | 0.871 |
| E619 | -0.962 | A668 | 0.047 | N717 | 0.016 | A766 | 0.029 | R815 | 0.827 |
| V620 | -0.134 | G669 | 0.004 | F718 | 0.006 | L767 | -0.006 | S816 | -0.221 |
| P621 | 0.165 | I670 | -0.015 | T719 | -0.014 | T768 | -0.053 | F817 | 0.021 |
| V622 | 0.032 | C671 | 0.062 | I720 | -0.024 | G769 | 0.040 | I818 | -0.014 |
| A623 | -0.071 | A672 | -0.077 | S721 | 0.027 | I770 | -0.024 | E819 | -0.669 |
| I624 | -0.012 | S673 | -0.054 | V722 | -0.024 | A771 | 0.035 | D820 | -0.739 |
| H625 | 0.023 | Y674 | -0.012 | T723 | 0.011 | V772 | -0.050 | L821 | -0.068 |
| A626 | -0.024 | Q675 | -0.061 | T724 | -0.044 | E773 | -0.967 | L822 | -0.043 |
| D627 | -0.918 | T676 | 0.098 | E725 | -0.945 | Q774 | 0.062 | F823 | 0.026 |
| Q628 | -0.017 | Q677 | -0.080 | I726 | -0.020 | D775 | -0.427 | N824 | 0.018 |
| L629 | -0.051 | T678 | -0.110 | L727 | -0.079 | K776 | 0.941 | K825 | 0.890 |
| T630 | -0.071 | N679 | 0.007 | P728 | 0.128 | N777 | -0.003 | V826 | 0.018 |
| P631 | 0.103 | S680 | -0.044 | V729 | -0.055 | T778 | -0.045 | T827 | -0.017 |
| T632 | -0.014 | P681 | 0.148 | S730 | 0.056 | Q779 | -0.009 | L828 | 0.022 |
| W633 | 2.009 | R682 | 0.911 | M731 | -0.083 | E780 | -0.987 | A829 | 0.012 |
| R634 | 1.033 | R683 | 0.971 | T732 | 0.022 | V781 | -0.014 | D830 | -0.934 |
| V635 | 0.009 | A684 | -0.030 | K733 | 0.481 | F782 | -0.039 | A831 | -0.099 |
| Y636 | -0.015 | R685 | 0.936 | T734 | -0.005 | A783 | -0.006 | G832 | 0.003 |
| S637 | 0.015 | S686 | 0.086 | S735 | -0.072 | Q784 | -0.075 | F833 | -0.023 |
| T638 | -0.112 | V687 | 0.054 | V736 | -0.016 | V785 | 0.005 | I834 | -0.993 |
| G639 | 0.022 | A688 | -0.037 | D737 | -0.790 | K786 | 0.975 | | |
| S640 | -0.031 | S689 | -0.006 | C738 | 0.004 | Q787 | -0.038 | | |

**Table S4:** List of PC value for each amino acids with their sequence number for WT HR1-CH.

| AA Seq No | PC | AA SeqNo | PC | AA SeqNo | PC | AA SeqNo | PC |
|---|---|---|---|---|---|---|---|
| K835 | 1.893 | G885 | 0.072 | Q935 | -0.065 | D985 | -0.909 |
| Q836 | 0.082 | W886 | 0.003 | D936 | -0.861 | K986 | 0.381 |
| Y837 | 0.002 | T887 | -0.087 | S937 | 0.022 | V987 | 0.074 |
| G838 | 0.025 | F888 | 0.025 | L938 | 0.058 | E988 | -0.867 |
| D839 | -0.939 | G889 | -0.007 | S939 | -0.022 | A989 | -0.066 |
| C840 | -0.054 | A890 | -0.061 | S940 | -0.138 | E990 | -0.396 |
| L841 | -0.022 | G891 | -0.001 | T941 | -0.110 | V991 | 0.032 |
| G842 | 0.003 | A892 | -0.011 | A942 | 0.054 | Q992 | 0.011 |
| D843 | -0.675 | A893 | 0.031 | S943 | 0.014 | I993 | -0.111 |
| I844 | -0.056 | L894 | -0.080 | A944 | 0.004 | D994 | -0.803 |
| A845 | -0.007 | Q895 | 0.095 | L945 | -0.012 | R995 | 0.991 |
| A846 | 0.013 | I896 | -0.117 | G946 | 0.044 | L996 | -0.017 |
| R847 | 0.672 | P897 | 0.079 | K947 | 1.051 | I997 | -0.016 |
| D848 | -0.905 | F898 | 0.047 | L948 | -0.067 | T998 | -0.117 |
| L849 | -0.017 | A899 | 0.074 | Q949 | -0.093 | G999 | 0.039 |
| I850 | 0.006 | M900 | -0.042 | D950 | -0.813 | R1000 | 0.789 |
| C851 | 0.061 | Q901 | -0.061 | V951 | 0.038 | L1001 | -0.016 |
| A852 | 0.024 | M902 | 0.024 | V952 | 0.002 | Q1002 | -0.007 |
| Q853 | 0.000 | A903 | 0.023 | N953 | -0.039 | S1003 | 0.084 |
| K854 | 0.787 | Y904 | 0.000 | Q954 | 0.088 | L1004 | -0.031 |
| F855 | 0.040 | R905 | 0.777 | N955 | 0.032 | Q1005 | -0.006 |
| N856 | -0.044 | F906 | -0.054 | A956 | 0.001 | T1006 | -0.054 |
| G857 | -0.024 | N907 | -0.036 | Q957 | -0.096 | Y1007 | 0.039 |
| L858 | -0.023 | G908 | 0.093 | A958 | 0.046 | V1008 | -0.040 |
| T859 | 0.047 | I909 | -0.049 | L959 | -0.055 | T1009 | 0.011 |
| V860 | 0.060 | G910 | -0.019 | N960 | 0.019 | Q1010 | 0.044 |
| L861 | -0.010 | V911 | -0.055 | T961 | -0.018 | Q1011 | 0.066 |
| P862 | 0.016 | T912 | 0.056 | L962 | 0.030 | L1012 | 0.026 |
| P863 | 0.104 | Q913 | -0.055 | V963 | 0.001 | I1013 | -0.004 |
| L864 | -0.024 | N914 | 0.068 | K964 | 0.874 | R1014 | 0.770 |
| L865 | -0.037 | V915 | -0.034 | Q965 | 0.085 | A1015 | -0.017 |
| T866 | -0.153 | L916 | -0.012 | L966 | -0.010 | A1016 | 0.037 |
| D867 | -0.912 | Y917 | 0.042 | S967 | -0.028 | E1017 | -0.943 |
| E868 | -0.745 | E918 | -0.954 | S968 | 0.064 | I1018 | -0.018 |
| M869 | 0.030 | N919 | -0.086 | N969 | -0.025 | R1019 | 0.857 |
| I870 | -0.060 | Q920 | 0.039 | F970 | 0.082 | A1020 | 0.021 |
| A871 | 0.034 | K921 | 1.050 | G971 | 0.071 | S1021 | -0.017 |
| Q872 | -0.005 | L922 | -0.014 | A972 | -0.020 | A1022 | -0.029 |
| Y873 | 0.011 | I923 | -0.013 | I973 | -0.071 | N1023 | 0.115 |
| T874 | 0.008 | A924 | 0.032 | S974 | -0.151 | L1024 | -0.029 |
| S875 | -0.017 | N925 | 0.035 | S975 | -0.033 | A1025 | 0.001 |
| A876 | 0.031 | Q926 | -0.016 | V976 | 0.016 | A1026 | 0.000 |
| L877 | 0.008 | F927 | -0.003 | L977 | 0.033 | T1027 | 0.016 |
| L878 | -0.027 | N928 | 0.029 | N978 | 0.035 | K1028 | 0.977 |
| A879 | -0.012 | S929 | -0.022 | D979 | -0.727 | M1029 | 0.008 |
| G880 | 0.035 | A930 | -0.029 | I980 | 0.079 | S1030 | -0.043 |
| T881 | -0.027 | I931 | -0.028 | L981 | 0.014 | E1031 | -0.990 |
| I882 | 0.018 | G932 | 0.056 | S982 | -0.006 | C1032 | -0.021 |
| T883 | -0.059 | K933 | 0.968 | R983 | 0.772 | V1033 | -0.070 |
| S884 | -0.075 | I934 | -0.025 | L984 | -0.047 | L1034 | -0.709 |

**Table S5:** NL bonding information for WT SD2-FP shown in **Figure 4(a)** and **4(b)**.

| **P681** | **AA1** | **AA2** | **BL** | **BO** |
|---|---|---|---|---|
| C-C | P681(C) | R683(CA) | 4.3112 | 0.0003 |
| C-O | P681(C) | N679(O) | 3.8317 | 0.0016 |
| C-O | A684(CA) | P681(O) | 3.5811 | 0.0015 |
| C-O | R683(C) | P681(O) | 3.9893 | 0.0012 |
| C-O | R685(CA) | P681(O) | 4.0486 | 0.0012 |
| C-O | A684(C) | P681(O) | 3.7966 | 0.0011 |
| C-O | P681(CA) | N679(O) | 3.9701 | 0.0009 |
| C-O | R683(CA) | P681(O) | 4.1355 | 0.0006 |
| C-O | P681(C) | R685(O) | 4.0265 | 0.0001 |
| H-C | R685(H) | P681(CA) | 4.3914 | 0.0005 |
| H-C | N679(HB2) | P681(CG) | 4.4225 | 0.0002 |
| H-C | A684(H) | P681(CA) | 3.7109 | 0.0001 |
| H-C | P681(HB2) | A688(C) | 4.1737 | 0.0001 |
| H-C | A688(HA) | P681(CA) | 4.3491 | 0.0001 |
| H-H | N679(HB2) | P681(HD3) | 3.6204 | 0.0001 |
| H-H | N679(HB2) | P681(HG2) | 4.1921 | 0.0001 |
| H-H | P681(HB2) | A688(HB1) | 4.4138 | 0.0001 |
| N...H | R683(N) | P681(HA) | 4.3121 | 0.0002 |
| N...H | P681(N) | N679(HB2) | 4.4207 | 0.0001 |
| N-C | R685(N) | P681(C) | 4.2318 | 0.0018 |
| O...H | A684(H) | P681(O) | 1.9853 | 0.0221 |
| O...H | R685(H) | P681(O) | 2.0272 | 0.0195 |
| O...H | A684(HB3) | P681(O) | 4.1600 | 0.0001 |
| **D614** | | | | |
| C-O | G593(C) | D614(OD1) | 3.8390 | 0.0008 |
| C-O | D614(CG) | F592(O) | 4.1658 | 0.0001 |
| C-O | D614(C | F592(O) | 4.4372 | 0.0001 |
| C-O | F592(C) | D614(OD1) | 4.4956 | 0.0001 |
| H-C | G593(HA3) | D614(CA) | 4.1604 | 0.0002 |
| H-C | N616(H) | D614(C) | 4.3890 | 0.0001 |
| H-C | D614(H) | G593(C) | 4.4405 | 0.0001 |
| N...H | D614(N) | G648(HA3) | 4.2644 | 0.0002 |
| N-C | N616(N) | D614(C) | 4.2832 | 0.0002 |
| N-O | G593(N) | D614(OD1) | 4.1450 | 0.0011 |
| O...H | G593(HA2) | D614(OD1) | 2.3697 | 0.0006 |
| O...H | G594(H) | D614(OD1) | 2.8758 | 0.0009 |
| O...H | N616(H) | D614(O) | 4.0740 | 0.0001 |
| O...H | D614(HB3) | F592(O) | 4.1371 | 0.0001 |

**Table S6:** NL bonding information for mutated R681 shown in **Figure 4(c)** and **4(d)**.

| **R681** | **AA1** | **AA2** | **BL** | **BO** |
|---|---|---|---|---|
| C-C | E654(CD) | R681(CZ) | 4.4034 | 0.0007 |
| C-C | N679(C) | R681(CA) | 4.3343 | 0.0002 |
| C-C | R681(C) | R683(CA) | 4.4096 | 0.0003 |
| C-O | R681(CZ) | E654(OE1) | 3.9477 | 0.0016 |
| C-O | R681(C) | N679(O) | 3.8731 | 0.0015 |
| C-O | R681(CA) | N679(O) | 3.9977 | 0.0007 |
| C-O | R683(C) | R681(O) | 4.1159 | 0.0009 |
| C-O | R683(CA) | R681(O) | 4.2376 | 0.0005 |
| C-O | A684(CA) | R681(O) | 3.6842 | 0.0012 |
| C-O | A684(C) | R681(O) | 3.8998 | 0.0008 |
| C-O | R685(CA) | R681(O) | 4.1073 | 0.0010 |
| H-C | R681(HH12) | E654(CG) | 3.9178 | 0.0050 |
| H-C | R681(HH11) | E654(CD) | 3.8786 | 0.0006 |
| H-C | R681(HB3) | A688(CB) | 4.0259 | 0.0002 |
| H-C | R681(HB2) | A688(C) | 4.0362 | 0.0001 |
| H-C | A684(H) | R681(CA) | 3.8994 | 0.0003 |
| H-C | A688(HB2) | R681(CG) | 4.4072 | 0.0002 |
| H-C | A688(HA) | R681(CA) | 4.0814 | 0.0001 |
| H-H | R681(HB3) | A688(HB2) | 3.6654 | 0.0002 |
| H-H | R681(HB2) | A688(HB1) | 4.2281 | 0.0002 |
| H-H | R681(HB3) | A688(HB3) | 4.1173 | 0.0001 |
| N...H | R683(N) | R681(HA) | 4.4977 | 0.0001 |
| N-C | R685(N) | R681(C) | 4.3002 | 0.0013 |
| N-O | R681(NH2) | E654(OE1) | 4.1715 | 0.0002 |
| N-O | R681(NH2) | E654(OE2) | 4.2740 | 0.0001 |
| O...H | R681(HH12) | E654(OE1) | 1.8646 | 0.0579 |
| O...H | R681(HH12) | E654(OE2) | 2.3609 | 0.0012 |
| O...H | R681(HH11) | A688(O) | 3.3227 | 0.0001 |
| O...H | R681(HH11) | E654(OE2) | 3.6317 | 0.0003 |
| O...H | R681(HB2) | A688(O) | 3.7363 | 0.0001 |
| O...H | A684(H) | R681(O) | 2.1127 | 0.0161 |
| O...H | A684(HB3) | R681(O) | 4.2097 | 0.0001 |
| O...H | R685(H) | R681(O) | 2.1173 | 0.0167 |
| **G614** | | | | |
| C-O | F592(C) | G614(O) | 4.3773 | 0.0001 |
| C-O | G614(CA) | Y612(O) | 4.4818 | 0.0002 |
| H-C | G614(H) | Y612(CA) | 4.1262 | 0.0002 |

**Table S7:** NL bonding information for double mutation R681 & G614 shown in **Figure 4(e)** and **4(f)**.

| **R681** | **AA1** | **AA2** | **BL** | **BO** |
|---|---|---|---|---|
| C-C | N679(C) | R681(CA) | 4.2988 | 0.0002 |
| C-C | R681(C) | R683(CA) | 4.4065 | 0.0002 |
| C-O | R681(C) | N679(O) | 3.7966 | 0.0018 |
| C-O | R681(CA) | N679(O) | 3.9376 | 0.0008 |
| C-O | R681(C) | R685(O) | 3.9700 | 0.0001 |
| C-O | R683(C) | R681(O) | 4.1225 | 0.0009 |
| C-O | R683(CA) | R681(O) | 4.2607 | 0.0004 |
| C-O | A684(CA) | R681(O) | 3.6620 | 0.0011 |
| C-O | A684(C) | R681(O) | 3.8577 | 0.0009 |
| C-O | R685(CA) | R681(O) | 4.0559 | 0.0011 |
| H-C | R681(HB3) | A688(CB) | 4.0901 | 0.0002 |
| H-C | R681(HB2) | A688(C) | 4.1631 | 0.0001 |
| H-C | R681(HH11) | E654(CD) | 4.2840 | 0.0001 |
| H-C | A684(H) | R681(CA) | 3.8290 | 0.0002 |
| H-C | R685(H) | R681(CA) | 4.4735 | 0.0005 |
| H-C | A688(HA) | R681(CA) | 4.1218 | 0.0001 |
| H-H | R681(HB3) | A688(HB2) | 3.7532 | 0.0001 |
| H-H | R681(HB3) | A688(HB3) | 4.1294 | 0.0001 |
| H-H | R681(HB2) | A688(HB1) | 4.3108 | 0.0001 |
| O…H | R681(HH12) | E654(OE1) | 2.7545 | 0.0025 |
| O…H | R681(HH12) | E654(OE2) | 3.0339 | 0.0002 |
| O…H | A684(H) | R681(O) | 2.1218 | 0.0149 |
| O…H | A684(HB1) | R681(O) | 2.8768 | 0.0001 |
| O…H | A684(HB3) | R681(O) | 4.1990 | 0.0001 |
| O…H | R685(H) | R681(O) | 2.0797 | 0.0188 |
| N-C | R685(N) | R681(C) | 4.2576 | 0.0015 |
| N...H | R683(N) | R681(HA) | 4.4463 | 0.0001 |
| **G614** | | | | |
| C-O | G614(CA) | Y612(O) | 4.4363 | 0.0003 |
| H-C | G614(H) | Y612(CA) | 4.1801 | 0.0002 |
| N-C | N616(N) | G614(C) | 4.4429 | 0.0001 |

**Table S8.** AABP of two HR1-CH models.

| **Models** | **Total AABP** | **NN AABP** | **Non-local AABP** | **AABP from HB** | **No. of NL AAs** | **Data notation** |
|---|---|---|---|---|---|---|
| WT D950 | 1.163 | 1.021 | 0.143 | 0.154 | 6 | D950-1.163-1.021-0.143-0.154-0 |
| Mutated N950 | 1.093 | 1.009 | 0.084 | 0.106 | 6 | N950-1.093-1.009-0.084-0.106-1 |

**Table S9:** Bonding information for WT and mutated HR1-CH shown in **Figure S7.**

| D950 | BL | BO | AA1 | AA2 | N950 | BL | BO | AA1 | AA2 |
|---|---|---|---|---|---|---|---|---|---|
| C-C | 4.4115 | 0.0007 | D950(C) | Q954(CG) | C-C | 4.3161 | 0.0002 | L948(C) | N950(CA) |
| C-C | 4.4565 | 0.0001 | D950(C) | N953(CA) | C-O | 3.9113 | 0.0011 | N950(C) | L948(O) |
| C-C | 4.4797 | 0.0001 | D950(C) | V952(C) | C-O | 3.9991 | 0.0006 | N950(CA) | L948(O) |
| C-O | 3.9614 | 0.0009 | D950(CA) | G946(O) | C-O | 4.0112 | 0.0005 | N950(CA) | G946(O) |
| C-O | 4.0222 | 0.0007 | D950(C) | L948(O) | C-O | 3.9725 | 0.0007 | N953(CG) | N950(OD1) |
| C-O | 4.1299 | 0.0005 | D950(CA) | L948(O) | C-O | 3.9971 | 0.0008 | N953(C) | N950(O) |
| C-O | 3.6276 | 0.0014 | N953(CG) | D950(OD1) | C-O | 4.0473 | 0.0002 | N953(CA) | N950(O) |
| C-O | 3.8914 | 0.0012 | N953(C) | D950(O) | C-O | 3.8267 | 0.0010 | Q954(CA) | N950(O) |
| C-O | 3.9653 | 0.0003 | N953(CA) | D950(O) | C-O | 4.1235 | 0.0002 | Q954(CD) | N950(O) |
| C-O | 3.6803 | 0.0015 | Q954(CA) | D950(O) | H-C | 3.8882 | 0.0003 | N950(HA) | N953(CA) |
| C-O | 4.0099 | 0.0002 | Q954(CD) | D950(O) | H-C | 4.1012 | 0.0002 | N950(HB2) | G946(C) |
| H-C | 3.8349 | 0.0003 | D950(HA) | N953(CA) | H-C | 4.3837 | 0.0001 | N950(HD21) | G946(C) |
| H-C | 4.0555 | 0.0001 | D950(HB2) | G946(C) | H-C | 3.8755 | 0.0001 | N953(HD22) | N950(CB) |
| H-C | 4.1004 | 0.0001 | D950(H) | G946(CA) | H-C | 3.9892 | 0.0002 | N953(HB2) | N950(CB) |
| H-C | 4.4805 | 0.0001 | D950(HB3) | Q954(CD) | H-C | 4.0192 | 0.0002 | N953(HB2) | N950(CG) |
| H-C | 3.7144 | 0.0017 | N953(HD22) | D950(CB) | H-C | 4.3440 | 0.0001 | N953(HD21) | N950(CG) |
| H-C | 4.4845 | 0.0003 | N953(HD22) | D950(C) | H-C | 3.9659 | 0.0004 | Q954(H) | N950(CA) |
| H-C | 3.7387 | 0.0001 | N953(HB2) | D950(CG) | H-C | 4.0778 | 0.0001 | Q954(HB2) | N950(C) |
| H-C | 3.8752 | 0.0001 | N953(HB2) | D950(CB) | H-H | 3.7496 | 0.0002 | N950(HA) | N953(HD21) |
| H-C | 3.8121 | 0.0002 | Q954(H) | D950(CA) | H-H | 3.8780 | 0.0003 | N950(HA) | N953(HB3) |
| H-C | 3.9420 | 0.0001 | Q954(HB2) | D950(C) | H-H | 3.9617 | 0.0001 | N950(HB3) | N953(HB2) |
| H-H | 3.7331 | 0.0003 | D950(HA) | N953(HD21) | H-H | 4.0594 | 0.0001 | N950(HD21) | N953(HD22) |
| H-H | 3.8226 | 0.0003 | D950(HA) | N953(HB3) | N...H | 4.0470 | 0.0001 | K947(N) | N950(HD22) |
| H-H | 4.3138 | 0.0001 | D950(HB3) | Q954(HE21) | N...H | 4.0439 | 0.0002 | N950(ND2) | N953(HD22) |
| N...H | 4.1160 | 0.0002 | K947(N) | D950(H) | N...H | 4.1611 | 0.0003 | N950(N) | N953(HB2) |
| N...H | 3.8694 | 0.0001 | D950(N) | N953(HD22) | N...H | 4.4219 | 0.0001 | N950(ND2) | G946(HA3) |
| N...H | 4.1080 | 0.0004 | D950(N) | N953(HB2) | N...H | 4.4693 | 0.0001 | V952(N) | N950(HA) |
| N...H | 4.4338 | 0.0002 | V952(N) | D950(HA) | N...H | 4.3346 | 0.0001 | Q954(N) | N950(HA) |
| N-C | 4.2093 | 0.0011 | D950(N) | G946(C) | N-C | 4.0100 | 0.0005 | N950(ND2) | G946(C) |
| N-C | 3.8942 | 0.0039 | N953(ND2) | D950(CG) | N-C | 3.9414 | 0.0001 | N953(ND2) | N950(CG) |
| N-C | 3.9615 | 0.0012 | Q954(N) | D950(C) | N-C | 4.0431 | 0.0004 | Q954(N) | N950(C) |
| N-N | 4.4244 | 0.0002 | L948(N) | D950(N) | N-N | 4.3688 | 0.0002 | L948(N) | N950(N) |
| N-N | 4.4000 | 0.0005 | D950(N) | V952(N) | N-N | 4.3199 | 0.0001 | N950(N) | N953(ND2) |
| N-N | 4.2600 | 0.0001 | D950(N) | N953(ND2) | N-N | 4.4405 | 0.0004 | N950(N) | V952(N) |
| O...H | 2.0770 | 0.0201 | D950(H) | G946(O) | O...H | 2.1235 | 0.0172 | N950(HD22) | G946(O) |
| O...H | 1.6805 | 0.0686 | N953(HD22) | D950(OD1) | O...H | 2.3166 | 0.0090 | N950(H) | G946(O) |
| O...H | 3.0025 | 0.0004 | N953(HB2) | D950(O) | O...H | 2.0216 | 0.0254 | N953(HD22) | N950(OD1) |
| O...H | 3.3303 | 0.0005 | N953(HD21) | D950(OD1) | O...H | 2.9768 | 0.0006 | N953(HB2) | N950(O) |
| O...H | 3.8586 | 0.0011 | N953(HD22) | D950(OD2) | O...H | 3.5009 | 0.0002 | N953(HD21) | N950(OD1) |
| O...H | 1.8985 | 0.0321 | Q954(H) | D950(O) | O...H | 2.0562 | 0.0213 | Q954(H) | N950(O) |
| O...H | 3.6536 | 0.0001 | Q954(HE22) | D950(O) | O...H | 3.6756 | 0.0001 | Q954(HE22) | N950(O) |
| O...H | 4.0865 | 0.0003 | Q954(HG2) | D950(O) | O...H | 4.1025 | 0.0002 | Q954(HG2) | N950(O) |
| O...H | 4.3762 | 0.0001 | Q954(HB3) | D950(O) | | | | | |
